# Metaproteomics reveals age-specific alterations of gut microbiome in hamsters with SARS-CoV-2 infection

**DOI:** 10.1101/2024.11.12.623292

**Authors:** Marybeth Creskey, Fabiola Silva Angulo, Qing Wu, Levi Tamming, Emily E F Fekete, Kai Cheng, Zhibin Ning, Angela Wang, Patrícia Brito Rodrigues, Vinícius de Rezende Rodovalho, Marco Aurélio Ramirez Vinolo, Daniel Figeys, Xuguang Li, Francois Trottein, Xu Zhang

## Abstract

<u>The gut microbiome’s pi</u>votal role in health and disease is well-established. SARS-CoV-2 infection often causes gastrointestinal symptoms and is associated with changes of the microbiome in both human and animal studies. While hamsters serve as important animal models for coronavirus research, there exists a notable void in the functional characterization of their microbiomes with metaproteomics. In this study, we present a workflow for analyzing the hamster gut microbiome, including a metagenomics-derived hamster gut microbial protein database and a data-independent acquisition metaproteomics method. Using this workflow, we identified 32419 protein groups from the fecal microbiomes of young and old hamsters infected with SARS-CoV-2. We showed age-specific changes in the expressions of microbiome functions and host proteins associated with microbiomes, providing further functional insight into the dysbiosis and aberrant cross-talks between the microbiome and host in SARS-CoV-2 infection. Altogether this study established and demonstrated the capability of metaproteomics for the study of hamster microbiomes.

## Introduction

The coronavirus disease 2019 (COVID-19) pandemic resulted in significant morbidity and mortality, causing severe social and economic disruption around the world. Infection with severe acute respiratory syndrome coronavirus 2 (SARS-CoV-2) is correlated with observable changes in the gut microbiome including a reduction in microbial diversity and alterations in the relative abundance of pathogenic bacterial species^1–5^. These microbiome alterations have been associated with disease severity and may play a role in immune dysregulation and systemic inflammation observed in COVID-19 patients^2, 6^. Functional characterization of these taxonomic changes in the gut microbiome during SARS-CoV-2 infections could provide insights into disease mechanisms and potential microbiome-directed therapeutic strategies for disease management.

The hamster has emerged as a prominent animal model for infectious diseases, including COVID-19, and is often preferred over mice due to several advantages, including (1) hamsters are outbred animals conferring more genetic diversity over mice, and (2) the infectious disease progression seen in hamsters are more comparable to that of humans^7^. For coronavirus disease, the hamster angiotensin-converting enzyme 2 (ACE2), the main cellular receptor mediating viral entry, binds more strongly with the spike protein than the mouse ACE2, which is consistent with observations that the hamster experiences mild to severe disease with quantifiable clinical signs, weight loss, viral shedding and lung pathology^8, 9^. Notably, recent studies have shown that the SARS-CoV-2 infection in hamsters is associated with alteration of gut microbial composition, similar to observations in human studies. Sencio et al. reported alteration of hamster gut microbiota composition along with SARS-CoV-2 infection in young-adult animals^10^. The infection and disease severity were associated with the increase of opportunistic pathogens such as *Enterobacteriaceae* and *Desulfovibrionaceae*, and decrease of short-chain fatty acid (SCFA) producing bacteria such as *Ruminococcaceae* and *Lachnospiraceae*. On the other hand, Seibert et al. showed that the middle-aged hamster microbiome infected with SARS-CoV-2 shares some similarities with that of critically ill COVID-19 patients^11^. A further study in high fat/high cholesterol diet induced obese nonalcoholic steatohepatitis (NASH) hamsters demonstrated that more severe disease activity was developed following SARS-CoV-2 infection in NASH hamsters with dysbiosis^12^. In the meantime, metabolomic analyses have also demonstrated the critical role of microbiota metabolites such as SCFA and deoxycholic acid in host resistance to SARS-CoV-2 infection and further host immune responses to infection, impacting disease severity^13, 14^.

However, despite its widespread use of hamster animal models, there remains a notable gap in the study of hamster gut microbiome with functional meta-omics methods, including metagenomics and metaproteomics. The majority of the current hamster gut microbiome studies were performed using the 16S rDNA amplicon sequencing approach. Currently, there are no published studies utilizing metaproteomics techniques in hamsters, and there are no hamster gut microbial gene/protein databases that are needed for metaproteomic identifications. To fill this knowledge gap, we constructed, evaluated and made publicly available hamster gut microbiome protein databases by combining both an in-house and a previously published shotgun metagenomic sequencing dataset. A remarkable advance of quantitative proteomics, including metaproteomics, in recent years is the wide application of data-independent acquisition mass spectrometry (DIA-MS)^15–18^. Therefore, this study utilized the DIA method, incorporating parallel accumulation serial fragmentation (PASEF)^19^, to develop a DIA-PASEF metaproteomics workflow for studying the hamster microbiome. We established a two-stage PASEF metaproteomic workflow, including a first stage data-dependent acquisition (DDA) PASEF analysis of pre-fractionated pooled samples for generating a tailored spectral library/database, and a second stage of DIA-PASEF analysis for efficiently identifying and quantifying both the gut microbial and host proteins in feces of hamsters.

As a proof of concept for the methodology, we analysed stool from young (2-month old) and old (22-month old) male Syrian golden hamsters infected with SARS-CoV-2 followed by sampling at day 0, 7, 15, 30, and 45. The established workflow enables in-depth profiling of the fecal microbial proteins, taxonomic compositions and functions of the hamster fecal microbiomes, and reveals distinct clustering patterns at all feature levels for old hamsters at day 7 after SARS-CoV-2 infection. The hamster gut microbial protein and spectral library databases, the DIA-PASEF metaproteomic workflow, and the deep metaproteomic dataset of SARS-CoV-2 infection provided in this study altogether are valuable resources for the microbiome study in hamster disease models.

## Results

### Constructing hamster gut microbial protein databases for metaproteomics

A protein sequence database is needed for shotgun proteomic or metaproteomic study to efficiently interpret mass spectra through peptide-spectrum matching (PSM). For the study of gut microbiomes, extensive metagenomic sequencing has been used to generate gut microbial gene catalogs or genome databases in hosts, such as humans and mice^20–22^. Previous study has shown limited overlap between the human and mouse gut microbial gene catalogs^21^, suggesting a need for host-specific gut microbial gene/protein databases for metaproteomic identification. Since there is currently no available hamster gut microbial gene catalog or genome database, we generated an in-house shotgun metagenomic sequencing dataset derived from 3 young (2-month old) and 6 old hamsters (22-month old), consisting of 434 million high quality sequencing reads. We used a previously established SqueezeMeta (v1.6.4) workflow^23^ for processing the metagenomic sequencing data. Briefly, high quality sequencing reads were first assembled using Megahit^24^, and open reading frame (ORF) or genes were predicted using Prodigal^25^ to generate a gene catalog database MGDB-V1, consisting of 1,730,340 genes or translated protein sequences (Figure 1A; details in method section). Taxonomic annotation and quantitation of the shotgun metagenomic sequencing data showed that, similar to human and mouse gut microbiota, Bacillota and Bacteroidota were the most abundant phyla in hamster gut microbiota. Additionally, *Thermodesulfobacteriota*, *Pseudomonadota*, *Deferribacterota* and *Actinomycetota* were among the top abundant phyla. The most abundant families include *Oscillospiraceae*, *Lachnospiraceae*, *Muribaculaceae*, *Rikenellaceae*, *Bacteroidaceae*, *Prevotellaceae* and *Desulfovibrionaceae* (Supplementary Table S1).

**Figure 1.**
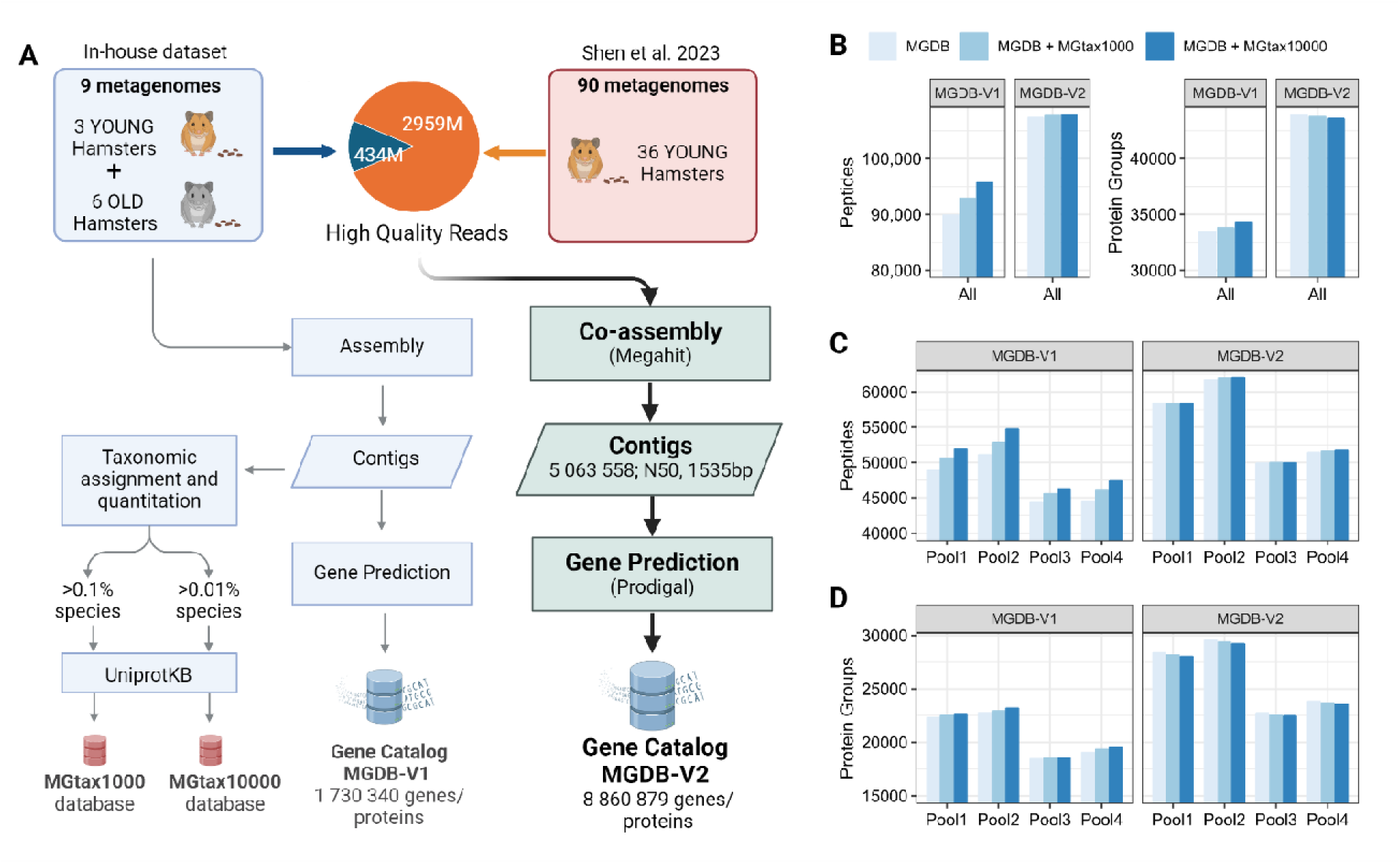
Hamster gut microbial gene catalog database construction and metaproteomic evaluation. (A) Gene catalog construction workflow using either in-house dataset only (MGDB-V1) or co-assembly (MGDB-V2) with a previous dataset by Shen et al. 2023^26^. Metagenomics based taxonomy derived proteome databases (MGtax) were established based on high abundant species and UnprotKB reference protein databases. (B-D) Different databases were used for search for 4 pooled sample metaproteomic data generated in this study. The numbers of identified peptides and protein groups for either all samples (B) or individual samples (C-D) are shown.

Usually the gene catalog database is sufficient for metaproteomics identification, however due to the limited sequencing depth of a single metagenomic dataset, the coverage of the gene catalog database may be suboptimal. To overcome this limitation, we applied two strategies to improve the established hamster gut microbial protein database. In the first method, we augmented the MGDB-V1 with downloaded UniprotKB proteome databases for the abundant microbial species based on metagenomics data (MGtax1000 for species abundance >0.1%, or MGtax10000 for abundance >0.01% according to contig taxonomic annotation and sum abundances for all samples in the in-house young/old hamster cohort; Figure 1A and Supplementary Table S2). In the second method, we downloaded a previously published dataset consisting of 90 metagenomes with 2959 million high quality sequencing reads from 30 young hamsters. This dataset was the only shotgun metagenomics dataset for hamster gut microbiome at the time of this project to the best of our knowledge. We therefore performed a co-assembly with both datasets to generate contigs and predict genes (Figure 1A). This co-assembly generated an updated version of the hamster gut microbial gene catalog database (MGDB-V2) with 8,860,879 proteins, over five times that of MGDB-V1.

We then used a fractionated DDA-PASEF dataset of hamster microbiome (i.e., 4 pooled samples with 8 fractionations per sample) to evaluate the performance of the different databases (Figure 1B). Notable increases in protein identifications were observed when MGtax1000 or MGtax10000 were added to the MGDB-V1 database (Figure 1C-D). Even higher numbers of peptides and protein groups were identified using MGDB-V2 than those with MGDB-V1 with or without MGtax augmentation. There was no obvious benefit observed for MGDB-V2 when augmenting with MGtax reference proteomes, indicating sufficient coverage of the database for MGDB-V2. We therefore selected MGDB-V2 without augmentation as the database for further DIA-MS metaproteomic workflow development.

### DIA-PASEF metaproteomics workflow to study hamster microbiome with SARS-CoV-2 infection

In this study, we aimed to demonstrate the capability of metaproteomics by studying the impacts of SARS-CoV-2 infection over time on hamster microbiomes. A total of 6 young and 6 old male Syrian hamsters were included, with one old hamster dying 24 days post-infection. Both groups experienced significant weight loss, with the lowest body weight recorded on Day 8 for the Old group and between Days 6 and 8 for the Young group (Supplementary Figure 1). While young hamsters began recovering quickly, nearly regaining their initial weight by Day 45, the older hamsters showed slower recovery and did not fully regain their initial body weight. In total, 58 fecal samples were collected spanning 5 time points (day 0, 7, 15, 30, 45) post infection for both groups (Figure 2). A standard fecal metaproteomic sample processing procedure was used, including a differential centrifugation to enrich microbial cells, protein extraction with sodium dodecyl sulfate (SDS) followed by SDS removal with acetone precipitation, and an in-solution trypsin digestion to obtain peptide samples for LC-MSMS analysis (Figure 2). To generate a reduced database and spectral library for DIA-PASEF data analysis, 4 pooled samples representing young, old, non-infected and infected animals, respectively, were generated, fractionated and analyzed with a DDA-PASEF mode on a timsTOF Pro2 MS. Each individual sample was analyzed with DIA-PASEF mode.

**Figure 2.**
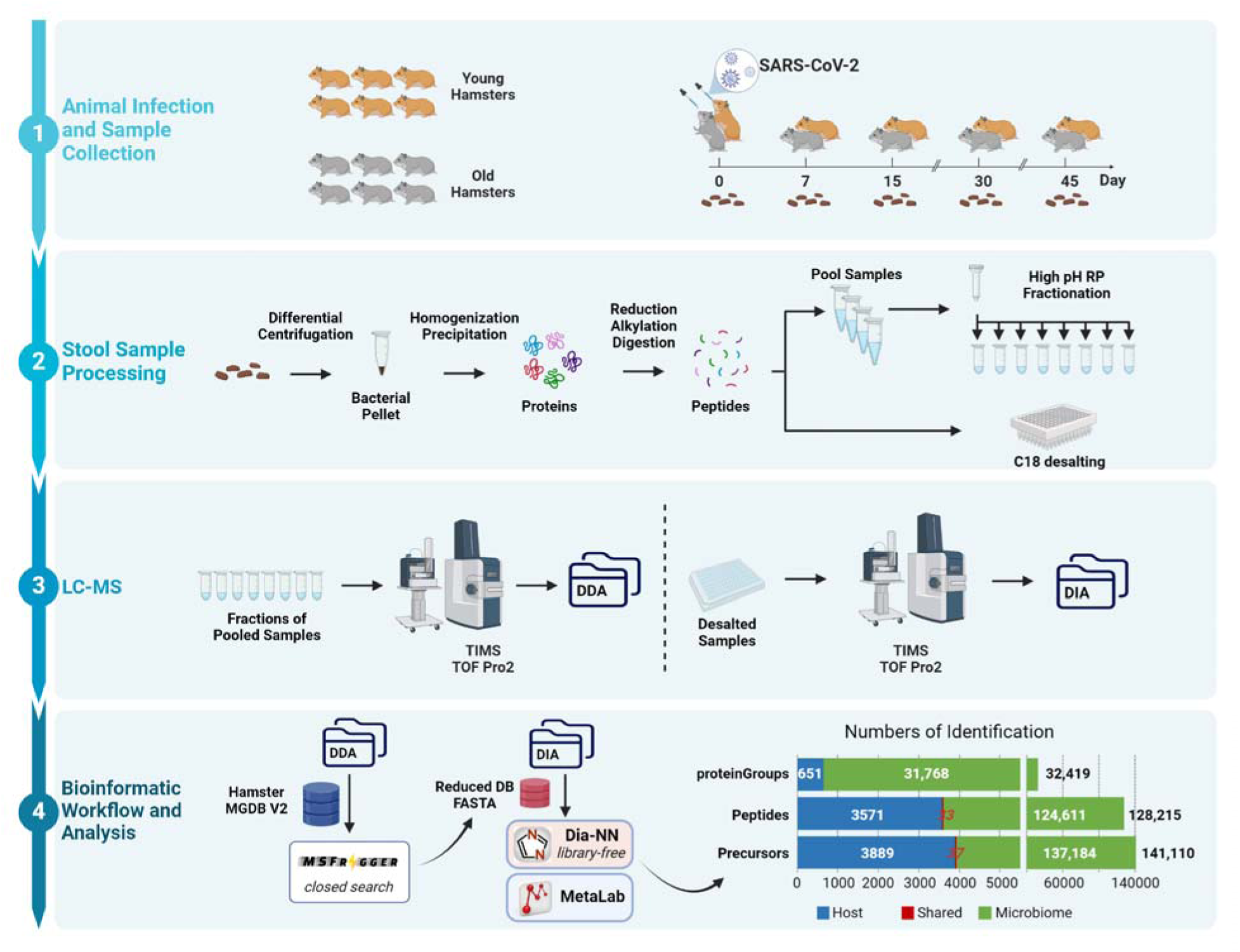
Experimental design and metaproteomics workflow for analyzing hamster microbiomes following SARS-CoV-2 infection. Animal experiment design, sample collection and processing steps are shown in the top panels (1 and 2). LC-MSMS analysis and bioinformatic workflow used in this study are shown in panels 3 and 4. Numbers of precursor, peptides and corresponding protein groups identified in the whole dataset are shown as well in panel 4.

We have previously reported that database reduction with DDA data using MSFragger followed by library-free search with DIA-NN performed the best for mouse metaproteomics data^18^. This finding was further validated here with hamster metaproteomics data (Supplementary Figure 2), and thereby we chose this workflow for the analysis of hamster metaproteomics data in this study. The host protein database was appended to the reduced .fasta database. In total, DIA metaproteomics identified 32,419 proteins and 128,216 peptides (Figure 2). Of these identifications, 651 were hamster proteins, and 31,768 were from the microbiome. Up to 57,852 precursors were identified per sample. Eight samples with precursor identification less than 30,000 (∼50% of the maximum identification; Supplementary Figure 3) were excluded from quantification and subsequent statistical analysis. Among all the identified protein groups, 11,627 were quantified in ≥70% samples (present in at least 35 out of the 50 samples). Non-supervised _≥_ principal component analysis (PCA) shows a time-based change after SARS-CoV-2 infection in both young and old hamsters (Figure 3A and Supplementary Figure 4). At Day 7, a perturbation of protein expression in both young and old animals is observed, with return to baseline at Day 15 and afterwards, representing acute phase at Day 7 and recovery phase after Day 15 (Figure 3A). When analysing host and microbiome proteins separately, obvious changes in host proteins were observed only in old hamsters at Day 7 following SARS-CoV-2 infection (Figure 3B). In contrast, marked impacts on microbiome proteins were observed in both young and old hamsters (Figure 3C). Interestingly, the old hamsters presented more extensive changes toward the same direction (2^nd^ principal component in PCA score plot) with young hamsters for the microbiome protein expressions, suggesting that the old animals tend to display more severe dysbiosis following infection. The latter may be associated with the increased disease severity often observed in elderly people^27^ as well as old hamsters^28–30^.

**Figure 3.**
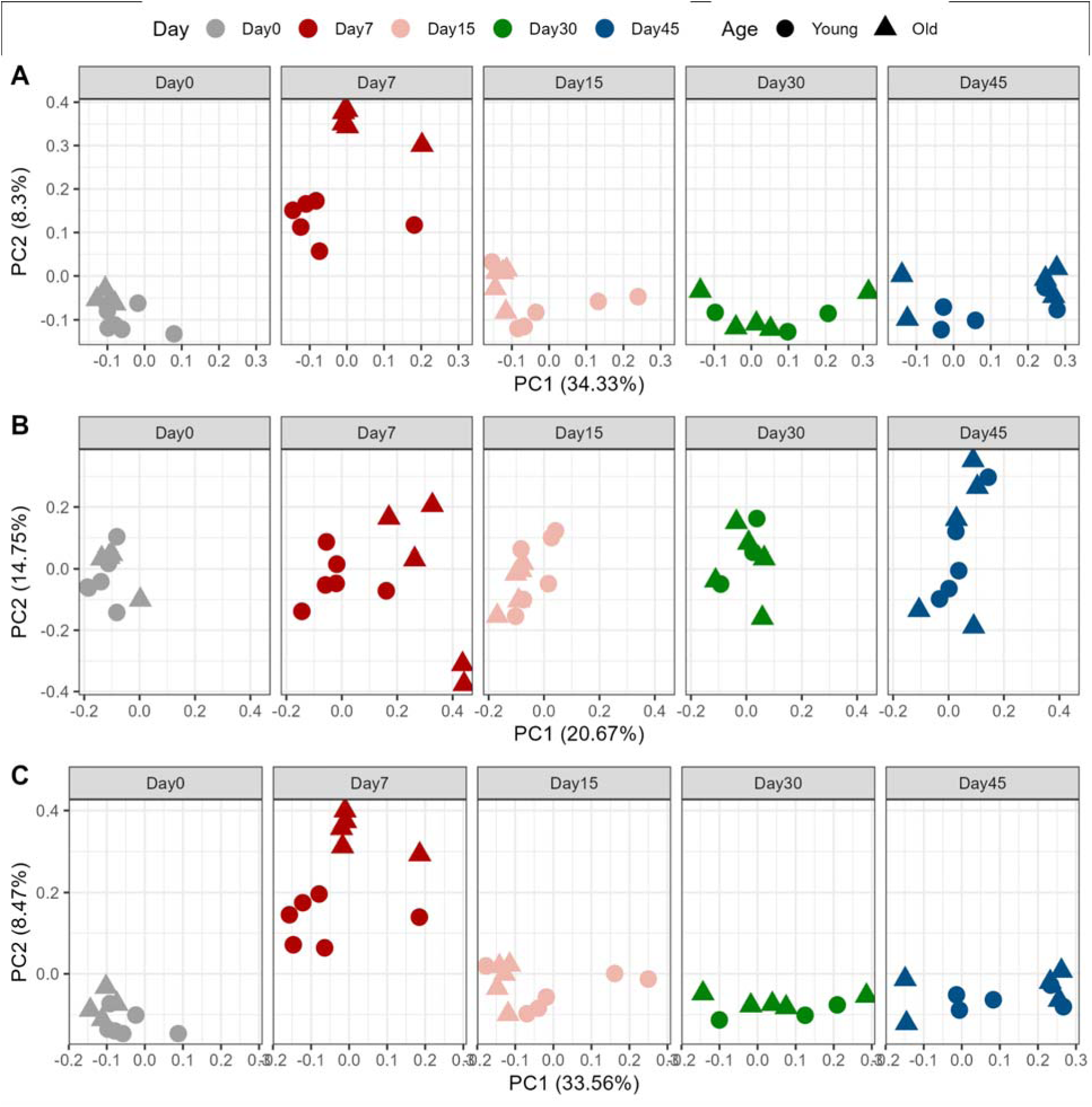
PCA score plots of proteins quantified in hamster feces following SARS-CoV-2 infection. PCA was performed using the normalized and log2-transformed protein intensities for all proteins (A), host proteins only (B) and microbiome proteins only (C), respectively. Only proteins that were quantified in >70% of the samples were used for analysis. PCA score plots were generated with R ggplot2 with facet according to time points.

### SARS-CoV-2 infection induced alteration of fecal host proteins in old hamsters

Metaproteomics has the advantage of quantifying both host and microbiome proteins at the same time. In this study, although a differential centrifugation process was included in the sample preparation to remove debris, large host cells, and proteins in the supernatant, we still identified 651 host proteins. These proteins may represent secreted host proteins closely associated with the microbial cell surface. To further examine the change of host proteins with SARS-CoV-2 infection, the relative abundance of host proteins for each sample was examined, by summing up the intensities of all hamster-derived proteins as a percentage of the total protein intensity. For young hamsters, a progressive increase in host protein amount is seen over the course of the study timeline, achieving significant differences at Day 30 and 45. A significant increase in host protein amount is observed for old hamsters on Day 7, but returns to baseline (Day 0) by Day 15 (Figure 4). An increasing trend is observed in old hamster from Day 15 to Day 45, resembling the pattern observed in young hamsters.

**Figure 4.**
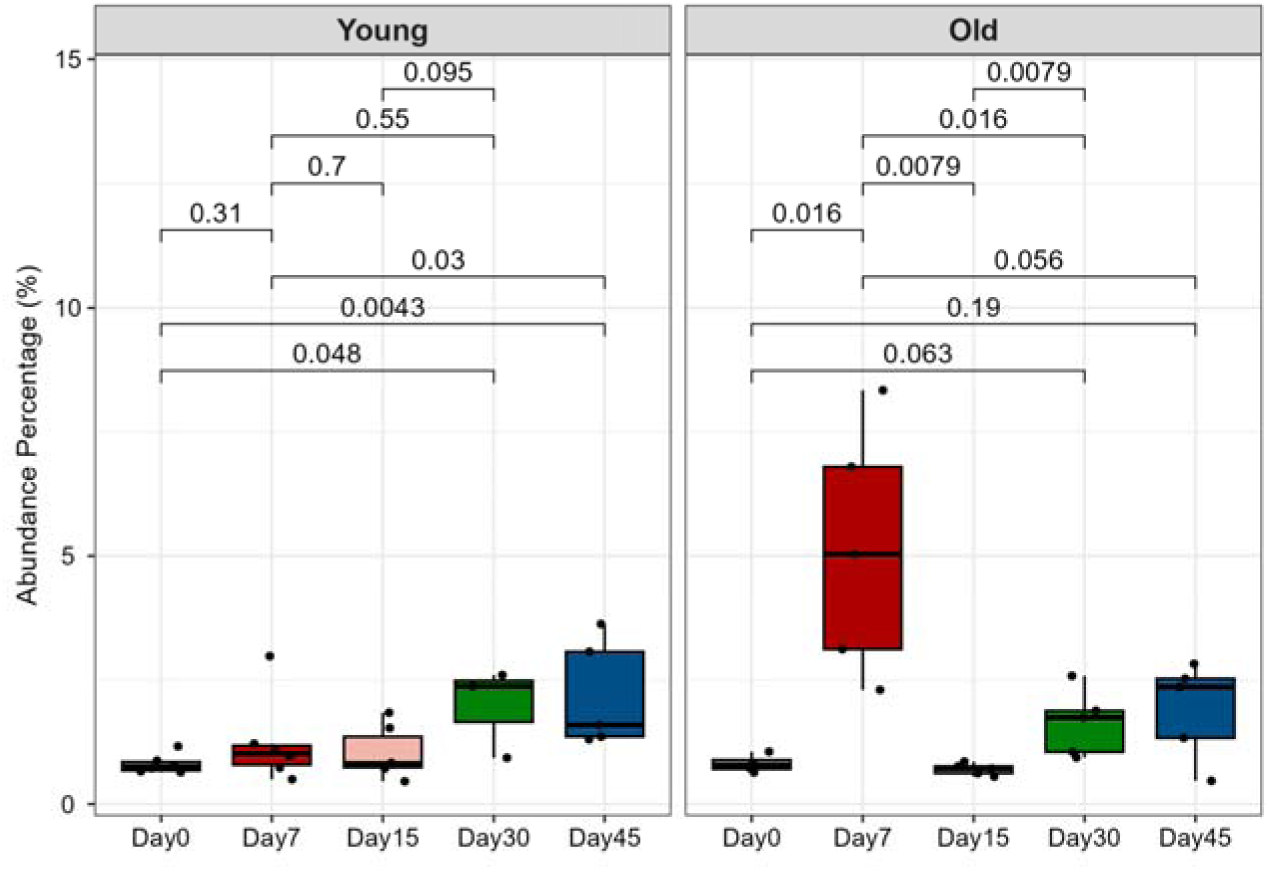
Relative abundances of host proteins in hamster fecal metaproteome. The percentage of sum intensities of all quantified host proteins in relation to the total intensity was calculated for each sample. Statistical significance (Wilcoxon test) and plotting were performed with ggpubr in R. P values are indicated for comparisons in both young and old groups for those with significance (P < 0.05) in either one group of hamsters.

We then sought to identify which host proteins were differentially regulated following SARS-CoV-2 infection. Since unsupervised PCA analysis showed that the most significant changes were at acute phase (Day 7) for both young and old groups (Figure 2 and Supplementary Figure 4), we next focused on identifying key proteins, microbial functions or taxa that drive the changes at acute phase for Young-Day7 or Old-Day7 groups compared to other samples. We performed partial least squares-discriminant analysis (PLS-DA) analysis for normalized host protein abundances, which achieved a model goodness of prediction (Q2) of 0.89 and the three clusters can be sufficiently separated with the first two PLS components (Supplementary Figure 5). Accordingly, 40 host proteins with variable importance projection (VIP) ≥1 in either PLS component 1 or 2 were selected (Figure 5 and Supplementary Table S3), among _≥_ which 13 were upregulated and 27 were down-regulated in the Old-Day7 group.

**Figure 5.**
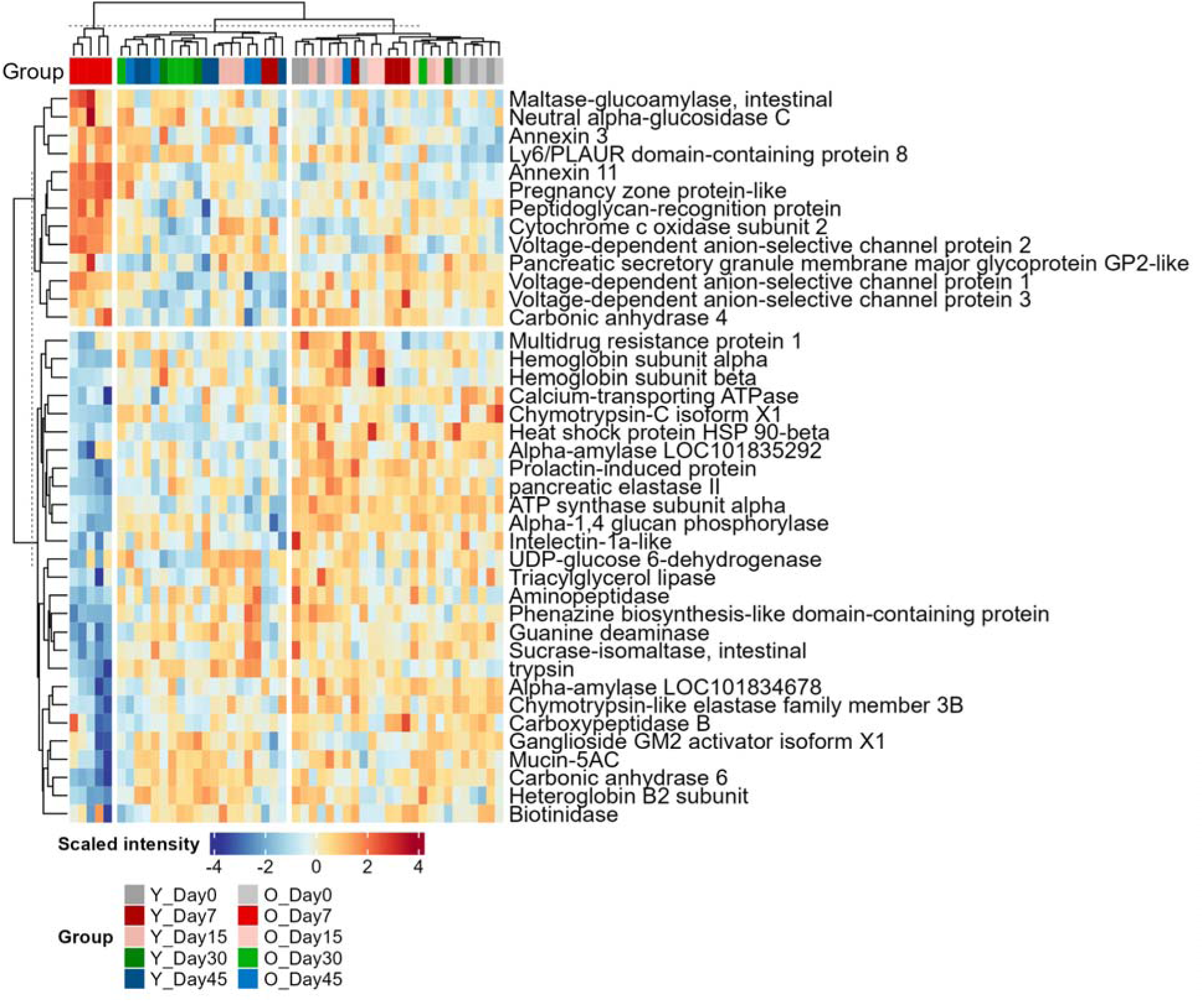
Heatmap of differentially abundant host proteins in Young-Day7 or Old-Day7 groups. Protein intensities were log2 transformed, scaled and are displayed as colours ranging from blue(low) to red(high) as shown in the key. Heatmap and clustering for both rows and columns are performed using the R ComplexHeatmap package.

The down-regulated host proteins in feces of SARS-CoV-2 infected hamsters include a diverse array of catabolic enzymes involved in carbohydrate metabolism (i.e., alpha-amylases, sucrase-isomaltase, alpha-1,4 glucan phosphorylase, and UDP-glucose 6-dehydrogenase), protein degradation (i.e., chymotrypsin-C, trypsin, elastase, chymotrypsin-like elastase family protein, aminopeptidase, and carboxypeptidase B), lipid metabolism and transport (i.e., triacylglycerol lipase and ganglioside GM2 activator isoform X1), as well as for metabolisms of nucleotide (guanine deaminase) and vitamins (biotinidase). Additionally, there are also decreases in proteins related with intestinal barrier function or immune responses to bacteria, such as mucin-5AC, intelectin-1a-like, prolactin-induced protein, and phenazine biosynthesis-like domain-containing protein. On the contrary, the up-regulated host proteins include those involved in recognizing extracellular components of Gram-positive and negative bacteria, such as peptidoglycan recognition protein, pancreatic secretory granule membrane major glycoprotein GP2-like protein, and Ly6/plaur domain containing protein 8. In addition, we also found the up-regulation of proteins related to mitochondria activity, including the voltage-dependent anion-selective channels (VDAC) proteins and Cytochrome c oxidase. Pregnancy zone protein-like is among the most significantly upregulated proteins in old hamsters with SARS-CoV-2 infection at Day 7, and has been reported to be potential biomarker for airway or mucosal infections in humans^31^. These alterations of host proteins associated with microbiomes may indicate an aberrant host-microbiome crosstalk during SARS-CoV-2 infection in hamsters.

The PLS-DA analysis for host protein data excluding Day 7 samples achieved a Q2 of 0.54 and demonstrated a gradual shift by time in recovery phase in PLS-DA score plot (Supplementary Figure 6). This observation aligns with the increasing total amount of host proteins in recovery phase (Day 15-45) in both young and old hamsters (Figure 4). A total of 48 proteins were identified with VIP threshold of 1 (Supplementary Figure 7 and Table S4). Among the 48 differentially abundant host proteins, eight presented increasing trend while 40 showed decreasing trend in particular in Day 30 and 45. We found that the abundances of several differentially expressed host proteins in acute phase were reversed in recovery phases, such as peptidoglycan recognition protein, VDACs, heteroglobin B2 and GP2-like proteins. However, some others such as Ly6/plaur domain containing 8 and Annexin 3 remain at high levels in recovery phase compared to Day 0 (Supplementary Figure 7), suggesting potential persistent impacts of viral infection on the intestinal epithelium function.

### SARS-CoV-2 infection induced alterations of microbiome functions in hamsters

We next assessed the impacts of SARS-CoV-2 infection on microbiome proteins and functions by analyzing the normalized microbial protein abundance data. To gain functional information, we annotated all the identified gut microbial proteins with GhostKOALA^32^ and calculated the abundance of each KEGG Orthology (KO) according to the annotation and protein abundances. In total, 22,158 (69.7%) out of the 31,768 microbial protein groups were annotated into 1571 KOs. Non-supervised PCA of the KO abundance data shows similar sample clustering to those with protein group abundances, namely an obvious shift only at Day 7 and with an age-dependent manner (Supplementary Figure 8).

We then utilized PLS-DA to identify differentially abundant KOs for acute phase in Young-Day7 and Old-Day7 groups compared to others. A Q2 of 0.93 and distinct separation of samples from Old-Day7, Young-Day7 and remainder clusters was achieved at the PLS component 1 (Supplementary Figure 9). Using a VIP threshold of 1, a total of 307 KOs were identified (Supplementary Table S5). Among the 307 differentially abundant KOs, 183 were enzymes and 52 were transporters, indicating significant alteration of microbial metabolism pathways induced by SARS-CoV-2 infections. Similarly, among the 58 KOs with VIP value >2, 31 were enzymes and 11 were transporters. There were 26 obviously up-regulated KOs and 32 down-regulated in the Old-Day7 group with the VIP threshold of 2. As can be seen in the heatmap (Figure 6), there was also a trend of shifting of Young-Day7 toward the Old-Day7 group direction, which agrees with the sample clustering in the PCA score plot. Among these KOs with VIP >2, six were directly associated with sulfide production and all of them were significantly upregulated in Old-Day7 and Young-Day7 groups. These KOs include dissimilatory sulfite reductases *Dsr* (K11180, K11181), sulfonate transport system permease protein (K15554), and three taurine/hypotaurine metabolism proteins to generate sulfite (K03851, K03852, K00259).

**Figure 6.**
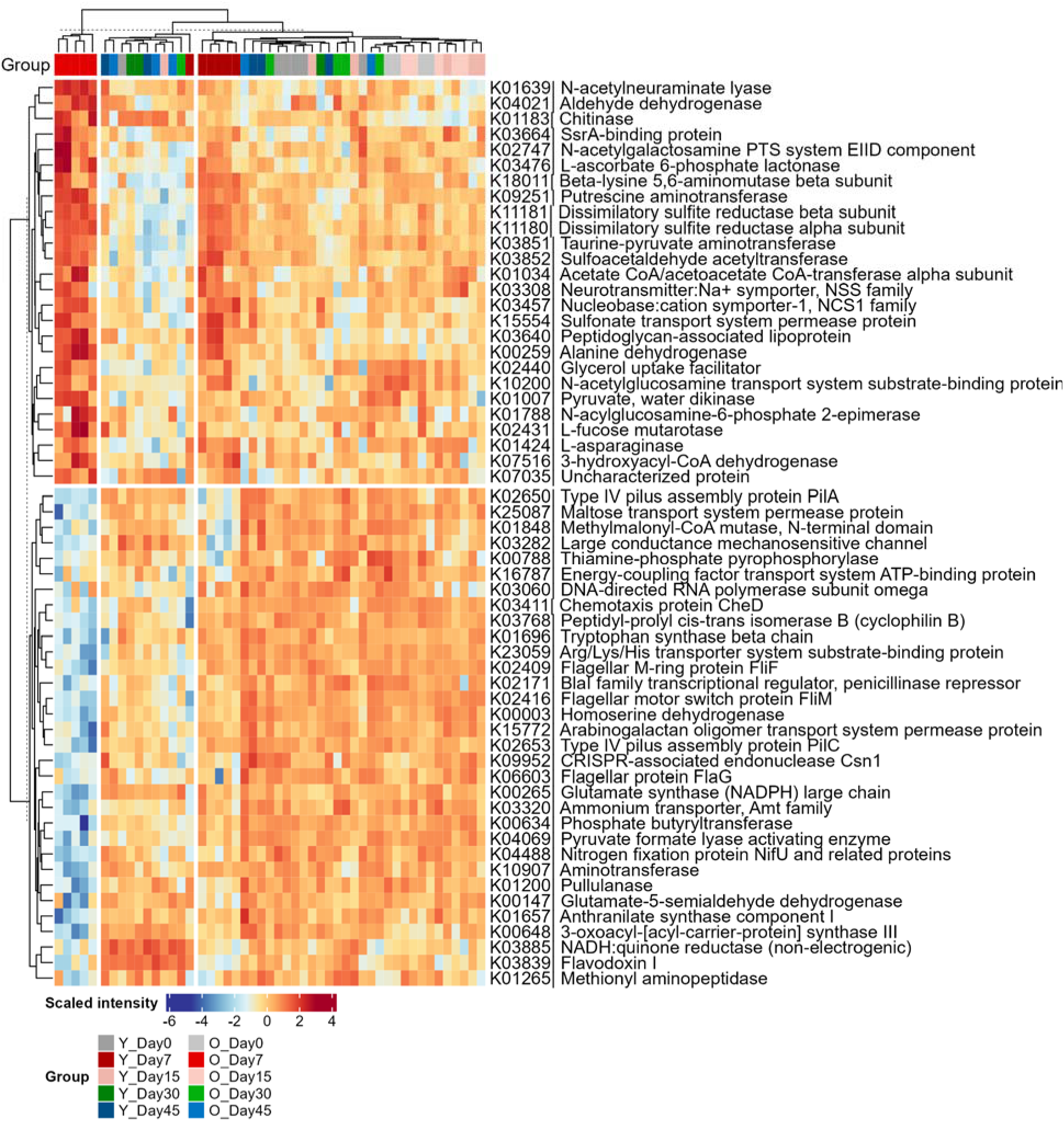
Heatmap of differentially abundant microbial functions. KEGG Orthology (KO) intensities were calculated by summing all protein intensities annotated to that KO. KO intensities are then log2 transformed, scaled and displayed as colours ranging from blue(low) to red(high) as shown in the key. Heatmap and clustering for both rows and columns are performed using the R ComplexHeatmap.

We also found an increase of peptidoglycan-associated lipoprotein (K03640), a key component of Gram-positive cell wall, in both Young-Day7 and Old-Day7 groups. This observation is in agreement with the upregulation of host peptidoglycan recognition protein in the Old-Day7 group observed in this study. It has been widely recognized that the gut microbiota has a pivotal role in neurological disorders through the gut-brain axis^33, 34^. We found that neurotransmitter:Na+ symporter-1 (NSS family, K03308) and putrescine aminotransferase (K09251) were upregulated in both Young-Day7 and Old-Day7 groups. Putrescine is one of the sources of inhibitory neurotransmitter gamma-aminobutyric acid (GABA)^35^, and putrescine aminotransferase is the first step of putrescine degradation. The upregulation of these proteins indicates that the gut microbiota may be involved in the alteration of neurotransmitter homeostasis in the gut of hamsters with SARS-CoV-2 infection. Additionally, we also found a significant change of vitamin degradation related functions, namely L-ascorbate 6-phosphate lactonase (K03476), being upregulated in COVID-19 hamsters, suggesting potential impacts of SARS-CoV-2 infection in shaping host intestinal homeostasis from various aspects.

A total of 32 KOs were downregulated (with VIP>2) at both Young-Day7 and Old-Day7 groups. In addition to the enzymes involved in amino acid and polysaccharide metabolisms, these downregulated KOs include two type-IV pilus assembly proteins PilA, PilC, three flagellar related proteins Flif, FliM, FliG, as well as chemotaxis protein CheD that is closely associated with flagellar motility. These observations suggest potential significant impacts of SARS-CoV-2 infection on the gut microbial cellular motility function in hamsters.

We next sought to identify microbial functions altered during the recovery phase of SARS-CoV-2 infection (Day 15-45). PLS-DA analysis, excluding Day 7 samples, yielded a Q2 of 0.66 and identified 45 KOs with VIP scores >2 (Supplementary Table S6 and Supplementary Figure 10). Heatmap of the differentially abundant microbial KOs revealed the most pronounced changes on Day 30 and Day 45 (Supplementary Figure 11). The majority of these significantly changed functions were associated with carbohydrate (sugars) transport and metabolism, alongside changes in amino acid metabolism (involving aspartate, glycine and glutamate) and ribosomal proteins. Additionally, a significant decrease in anaerobic sulfite reductases (K16950 and K16951) was observed at Day 30 and 45, despite no changes at Day 7 and 15 timepoints (Supplementary Figure 11).

### Metaproteomics revealed extensive taxonomic alterations in COVID-19 hamsters

This study identified 128,217 peptides with 66,997 of them being mapped to unique taxa, which were then used for taxonomic analysis using a peptide-centric workflow in MetaLab^36^. By using a threshold of minimum 3 distinctive peptides, all four superkingdoms can be identified with 62 phyla, 109 classes, 183 orders, 264 families, 401 genera and 419 species (Figure 7A and Supplementary Table S7). Here we focus on microorganisms, so we removed all sequences assigned to Eukaryota (except for Fungi) and calculated relative abundances at each taxonomic rank level. Similar to gut microbiomes of other mammals, Bacilota and Bacteroidota are the main bacterial phyla in hamster fecal microbiome (Figure 7B). An obvious increase of Bacteroidota and Pseudomonadota and decrease of Bacillota can be observed in the Old-Day7 group. These shifts persisted in the recovery phase, particularly on Days 30 and 45, in both young and old groups. These findings are in agreement with observations with shotgun metagenomic sequencing analysis of microbiomes in an independent hamster cohort study^28^. Family level analysis also showed that *Bacteroidaceae* and *Tannerellaceae*, the two abundant Bacteroidota families, are the most obviously increased in the Old-Day7 group compared to others (Supplementary Figure 12). We also observed a significant decrease of species diversity, richness as well as evenness in the Old-Day7 group compared to Day 0, while no significant difference was observed for young hamsters (Supplementary Figure 13).

**Figure 7.**
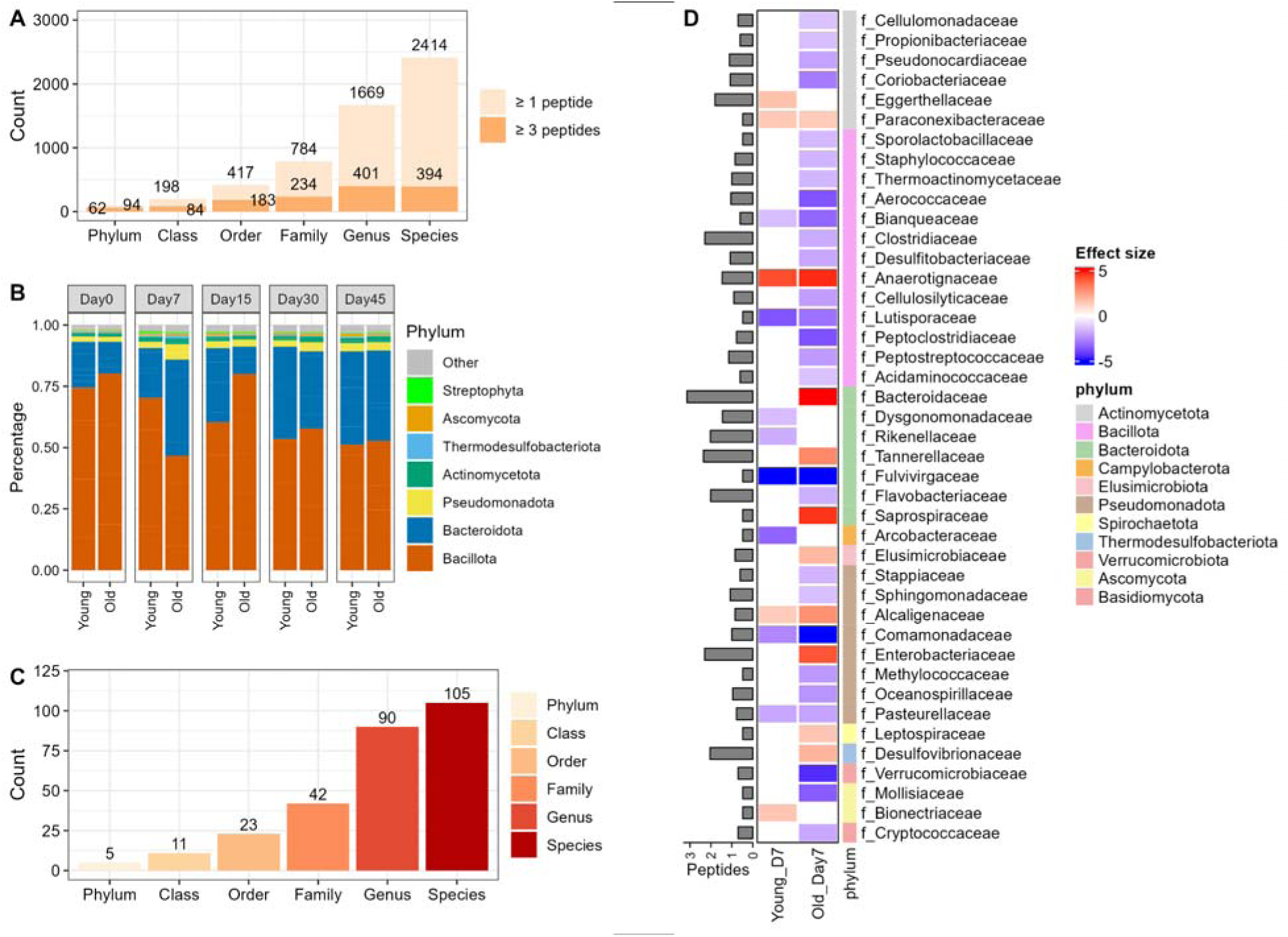
Metaproteome-based taxonomic composition and alterations in hamsters with SARS-CoV-2 infection. (A) Number of taxa identified at each rank level with a minimum of 1 or 3 distinctive peptides. Taxa with a minimum of 3 distinctive peptides were kept for further analysis in this study. (B) Phylum level composition for each group; (C) Numbers of significantly changed taxa at each rank level identified using MaAsLin2; (D) Significantly changed families identified with MaAsLin2. The heatmap shows the effect size with colours ranging from blue (down-regulated) to red (up-regulated) as shown in the key. Left side annotation bar plot shows the number (log10-transformed) of distinctive peptides for each family, while the right-side annotation shows the phylum assignment for each family as shown in the phylum legend.

To identify statistically significant differences of taxa at acute phase, we performed MaAsLin2 analysis^37^ based on a general linear model comparing Young-Day7 and Old-Day7 with the other groups. In total, 277 taxa were identified as significantly changed in either Young-Day7 or Old-Day7 group, including 5 phyla (Elusimicrobiota, Basidiomycota, Pseudomonadota, Thermodesulfobacteriota and Thermoproteota), 11 classes, 23 orders, 42 families, 90 genera and 105 species (Figure 7C and Supplementary Table S8). As shown in Figure 7D, the majority of the significantly different families were observed for the Old-Day7 group. Eight families showed significant changes in both groups, and all of them exhibited the same changing direction of changes in both young and old groups. *Fulvivirgaceae* was the most significantly down-regulated in both infection groups, while *Anaerotignaceae* (including *Anaerotignum faecicola* and *A. lactatifermentans*) was the most significantly up-regulated family in both groups. *Bacteroidaceae* is among the most significantly changed family and has the highest number of distinctive peptides (n=1396). Eleven species from *Bacteroidaceae* were identified as significantly changed in the Old-Day7 group and only two were decreased. *Tannerellaceae* (including *Parabacteroides distasonis* and *P. goldsteinii*), *Enterobacteriaceae* (mainly *Escherichia coli*, belong to Pseudomonadota) and *Desulfovibrionaceae* (mainly *Candidatus Bilophila faecipullorum, Bilophila wadsworthia, Mailhella massiliensis and Desulfovibrio legallii*) are also significantly increased in the Old-Day7 group and have high numbers of distinctive peptides (228, 199, and 110, respectively). Interestingly, all the four *Desulfovibrionaceae* species were significantly up-regulated in the Old-Day7 group, and three of them (except for *D. legallii*) were up-regulated in the Young-Day7 group as well. *Desulfovibrionaceae* species are known sulfate-reducing bacteria^38^. These observations are in agreement with the findings that KEGG functions related to sulfite reduction were up-regulated in both Young-Day7 and Old-Day7 groups.

Additionally, we also identified three fungal families that are significantly changed in either the Young-Day7 or Old-Day7 group, including *Mollisiaceae* and *Cryptococcaceae* that were down-regulated, and *Bionectriaceae* that are up-regulated (Figure 7D).

MaAsLin2 analysis comparing each individual group in the recovery phase to their corresponding baseline group (Day 0) identified 64 significantly changed taxa in young hamsters and 47 in old hamsters across Day 15, 30 or 45, using an adjusted P-value threshold of 0.05 (Supplementary Table S9). In the young group, the most significantly changed taxa included a decrease of *Ligilactobacillus* (Day 15), *Lachnospira* and *Roseburia inulinivorans* (Day 45), and an increase of *Bacteroides fragilis* (Day 45). In the old group, the most significant changes included an increase of *Ustilaginaceae* (belong to fungal phylum Basidiomycota, Day 30), and a decrease of *Coprococcus catus* and *Yeguia homins* (Day 30). Additionally, in young hamsters, significant increases were observed in the kingdom Fungi as well as fungal phyla Ascomycota and Basidiomycota on Days 15 and 45, along with a notable increase in the bacterial phylum Pseudomonadota and an archaeal phylum Euryarchaeota on Day 45.

### Co-occurrence analysis reveals taxon-specific functional alterations induced by SARS-CoV-2 infection

To examine the relationship between the host, microbial taxa and function alterations at acute phase, we performed a correlation analysis which identified 115 correlations with an absolute Spearman’s correlation coefficient (|r|) greater than 0.7 and an adjusted p-value threshold of 0.05. The most positive correlation is between *Tannerellaceae* and *Bacteroidaceae* (r = 0.91, p = 0), and the most negative correlation is between *Bacteroidaceae* and ATP5A1 (r = 0.86, p = 2.66E-15). With these significant correlations, three sub-networks were established (Figure 8A). The largest network is Network A, where *Tannerellaceae*, *Bacteroidaceae*, *Enterobacteriaceae*, and *Elusimicrobiaceae* were negatively correlated with a cluster of co-occurring down-regulated microbial functions, taxa and host proteins. In particular, the microbial cellular motility function related KOs (K03411, K02409, K02416) are positively correlated with each other and negatively correlated with *Tannerellaceae* and *Bacteroidaceae*.

**Figure 8.**
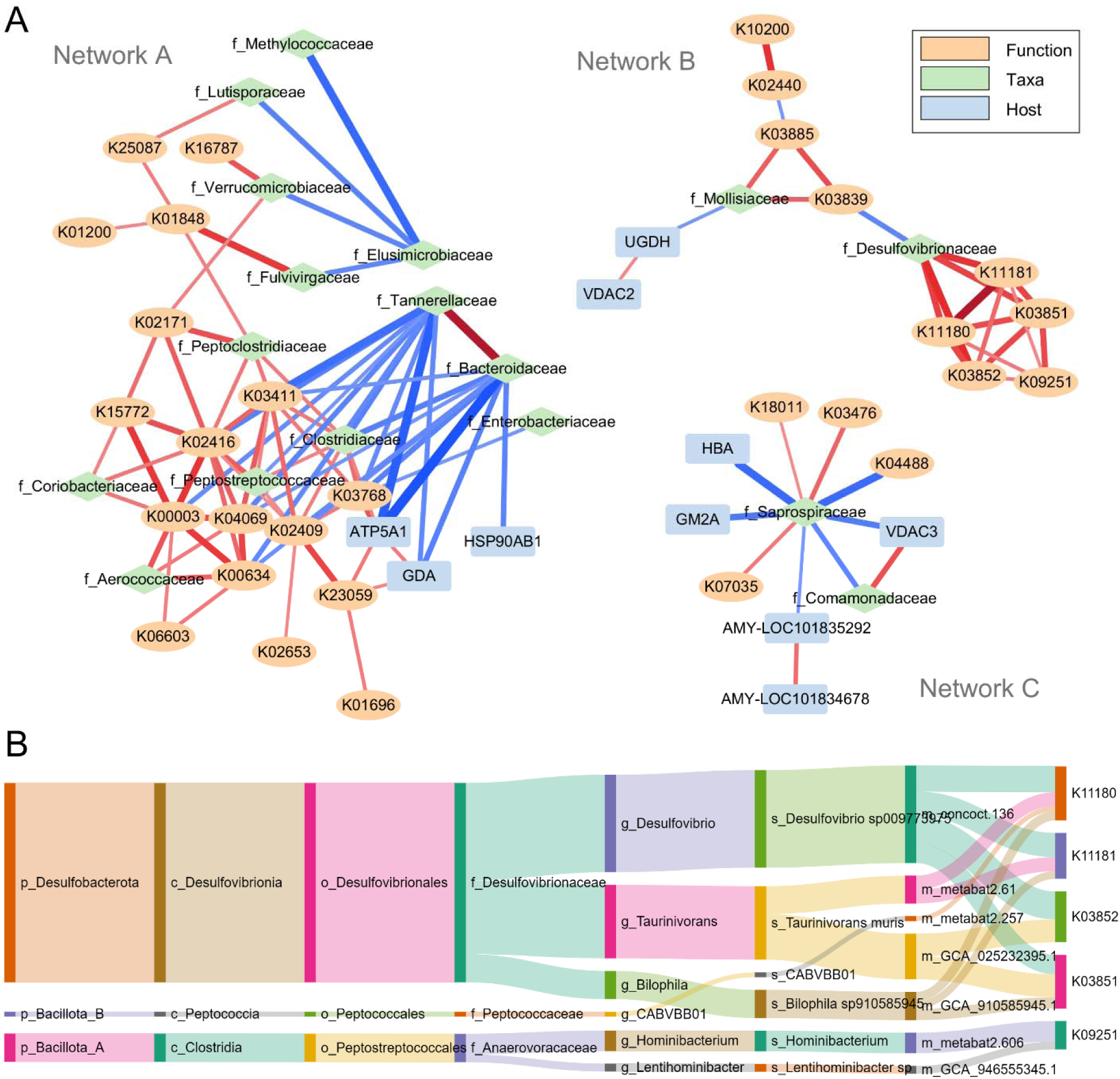
Correlation network of identified differentially abundant host proteins, microbial functions and taxa. (A) Pairwise Spearman correlations are calculated, and only correlations with a correlation |r| >0.7 and an adjusted P < 0.05 were used for network visualization using Cytoscape. Three sub-networks with > 4 nodes were shown. Color and shape of the nodes show host protein, microbial taxa or function type. The color and thickness of edges shows the direction (red as positive correlation, while blue as negative correlation) and coefficient (|r|) of correlation, respectively. (B) Sankey plot showing the OTF links between genomes, their taxonomic lineage and KEGG KOs positively associated with *Desulfovibrionaceae* in Network B.

The Network B is characterized by a close co-occurrence of the sulfate reducing bacteria *Desulfovibrionaceae* and KOs related to sulfite reduction (K11180 and K11181), taurine metabolism (K03851 and K03852), and putrescine aminotransferase (K09251). As mentioned above, putrescine is an important source of GABA, in both bacteria and mammalian cells^35^. The correlation between sulfate reducing bacteria and function with putrescine aminotransferase may indicate a role of sulfate reducing microbiota species in aberrant gut-brain axis homeostasis in SARS-CoV-2 infection. The hub of Network C is *Saprospiraceae*, which is among the most significantly up-regulated families in Old-Day7 hamsters and is worth further investigation.

To further explore the relationship between *Desulfovibrionaceae* and KOs related to sulfate metabolism, we then performed a taxon-function association analysis using MeteX at genome level^39^. The latter establishes associations between taxa and functions in metaproteomics based on the operational taxon-function unit (OTF) that is derived from linked peptides. By searching all OTFs with the 5 *Desulfovibrionaceae-*associated KOs in Network B, we showed that the four KOs related to sulfite reduction and taurine metabolism are derived from 5 genomes, 4 of which belong to *Desulfovibrionaceae* (Figure 8B). K09251 is uniquely associated with genome metabat2.606, a strain from genus *Hominibacterium*. The genome concoct.136 (*Desulfovibrio* sp.) expresses all four KOs related to sulfur metabolism, and the genome GCA_910585945.1 (*Bilophila* sp.) expresses 3 KOs except for K03852. Interestingly, the two genomes belonging to *Taurinivorans muris* express different functions, with GCA025232395.1 is only associated with taurine metabolism KOs, while metabat2.61 only with sulfite reduction KOs. By examining the relative abundance of each OTF, we observed a consistent increase for all the selected OTFs in the Old-Day7 group (Supplementary Figure 14). These observations indicate potential inter- and intra-species strain interactions in sulfur metabolism that may be involved in SARS-CoV-2 induced dysbiosis of hamster microbiome functions.

## Discussion

Although the COVID-19 pandemic has subsided, the disease is expected to persist as an endemic virus with seasonal surges, similar to influenza. Ongoing research into the disease’s mechanisms, vaccine efficacy, and long-COVID will continue to rely on hamsters as key model organisms. In addition to coronavirus diseases, hamsters have also been considered as a preferred small animal model for pathological and vaccine research for many emerging and re-emerging infectious diseases due to its ability to better meet regulatory guidelines for an appropriate animal model^7^. Previous hamster microbiome analysis primarily with 16S rDNA sequencing has demonstrated compositional changes that are associated with SARS-CoV-2 infection^10–12^. However, there is no hamster fecal metaproteomic study to date to the best of our knowledge. The lack of a hamster microbiome reference protein database is among the first challenges hampering the application of metaproteomics in hamster studies. In the current study, we have established a gene catalog as well as genome databases for the hamster gut microbiome by using an in-house as well as a published shotgun metagenomic sequencing dataset of both young and old hamsters. We also reported a comprehensive metaproteomic workflow with advanced DIA-MS methods, a multi-step database search strategy, and comprehensive downstream data analysis, enabling an in-depth functional characterization of hamster microbiomes. The application to a time course study of microbiomes in young and old hamsters with SARS-CoV-2 infection offers valuable insights into the dysbiosis and aberrant crosstalk between the microbiome and host, implicating elevated host antibacterial functions, opportunistic pathogen colonization, metabolic functions and their potential roles in impaired gut-brain homeostasis.

A major challenge for studying microbiomes using proteomics is the accurate and efficient identification of microbiome proteins^40^. This is particularly challenging for model organisms that are not well characterized for their microbiomes. Most of the current bioinformatic workflows for metaproteomic analysis are based on gene and/or genome catalogues that are derived from massive shotgun metagenomic sequencing. Increasing numbers of gene catalog or genome databases have been made available along with the community efforts. For example, EMBL-EBI MGnify has made available 12 comprehensive gut microbial genome catalogue databases for common model and non-model organisms^20^. These are invaluable reference resources enabling post-genomic applications, including metaproteomics. Unfortunately, there is so far no hamster microbiome gene catalog or genome databases available, despite the critical importance of hamster animal model and the gut microbiomes in COVID19 research. Here we addressed this knowledge gap by establishing a hamster gut microbial reference database, including 8.86 million genes/proteins and 926 genomes, through co-assembly of both young and old hamster, in-house and published shotgun metagenomic data. The size of the gene catalog database established here is close to the well-established integrated human gut microbial gene catalog (IGC, with 9.9 million non-redundant genes^22^). We demonstrated that the established gene catalog database provided sufficient coverage for metaproteomics protein identification, achieving >30,000 protein groups and >120,000 peptides with up to 57,000 peptides per sample. This depth of identification is also equivalent to the deepest metaproteomics application studies of human^17^ or mouse^41^ microbiomes so far.

We then performed the very first hamster metaproteomic study and examined the impacts of SARS-CoV-2 infection on the microbiome functionality in young and old hamsters over 45 days post infection (covering acute and recovery phase of diseases). Interestingly we found that the hamster metaproteome significantly changed at Day 7 but by Day 15 mostly returned to baseline. This observation aligns well with the fact that the phenotype recovery and viral clearance are completed 2∼3 weeks after infection in both hamsters and humans^2, 28^. More interestingly, the current metaproteomics study also demonstrated that the microbiome responses in young and old hamsters were not the same, with a notably higher extent of changes in old hamsters than those of young animals. These observations also agree with the shotgun metagenomics as well as the plasma metabolomics analyses for samples collected in an independent cohort of hamsters^28^. These age-specific alterations include significant decrease of microbial diversity only for Old-Day7 group (Supplementary Figure 13), more taxa that are significantly changed in the Old-Day7 group than the Young-Day7 group (Figure 7), as well as more extent of abundance alterations of key microbial functions (Figure 6). The observations in this hamster metaproteomic study align well with the more severe microbiome dysbiosis in elderly as well^42^. In humans, the elderly population is more severely impacted by COVID-19 due to age-related changes in metabolism and immune function. This demographic typically experiences high levels of inflammation, stress, catabolism, and increased energy and protein needs in their gastrointestinal tract^43^, and thereby lead to more vulnerable microbiomes.

One advantage of metaproteomics over genomic approaches is that it permits simultaneous identification and quantification of both host and microbiome proteins. This advantage enables the exploration of the crosstalk between the microbiome and the host immune system as a result of SARS-CoV-2 infection. We showed that while there is a general increasing trend of microbiome cell-associated host proteins in both young and old hamsters after SARS-CoV-2 infection, remarkable elevation of host mucosal protein secretion into the gut was observed in the Old-Day7 group (Figure 4). SARS-CoV-2 infects and replicates within enterocytes in the small intestine, specifically targeting the intestinal mucosa^44^. A healthy mucus layer is essential for maintaining intestinal homeostasis by supporting the symbiotic relationship between the host and gut microbiota. This mucus layer not only provides spatial separation between microbes and the intestinal epithelium but also acts as a selective filter, facilitating crucial host-microbe interactions. Further looking at the significantly altered host proteins, this study demonstrated a wide spectrum of down-regulation of catabolic enzymes and barrier function related proteins, while up-regulation of antibacterial and mitochondria activity related proteins. For example, we found the up-regulation of Ly6/plaur domain-containing protein 8 and peptidoglycan recognition protein in the Old-Day7 group, both of which are bacterial extracellular component binding proteins and play key roles in maintaining the mucosal barrier by preventing the invasion of bacteria into the inner mucus layer of the colon epithelium^45^. These observations of altered host protein secretion into the gut indicate significant disruption of intestinal homeostasis in old hamsters with SARS-CoV-2 infection.

Metaproteomics can provide biomass-based taxonomic compositions as well as taxon-specific protein or functional expressions. In this study, with a metaproteome-based taxonomics analysis we see a decrease in microbial diversity and an increase in opportunistic pathogens *Enterobacteriaceae* (mainly *Escherichia coli*), families in Bacteroidota, and sulfate-reducing *Desulfovibrionaceae* in Old-Day7 hamsters with SARS-CoV-2 infection. These taxonomic alterations also align with those of human patients with COVID-19 which showed significantly decreased bacterial diversity with enrichment of opportunistic pathogens, such as *Streptococcus*, *Veillonella*, *Fusobacterium* and *Escherichia* ^1^. In addition, the elevation of sulfate-reducing bacteria has also been reported to be implicated in many diseases, including inflammatory bowel diseases. Mottawea et al. demonstrated that a bacterium *Atopobium parvulum* in the gut of pediatric IBD patients can produce H_2_S, which lead to the onset of colitis; while administering H_2_S scavenger can mitigate *A. parvulum*-induced colitis in animal models^46^. Depletion of sulfate-reducing *Desulfovibrionaceae* is also commonly associated with beneficial effects of dietary intervention for alleviating metabolic disorders in humans^47^. By using a co-occurrence analysis as well as taxon-function linking with MetaX^39^, the current study demonstrated the upregulation of *Desulfovibrionaceae* specific dissimilatory sulfite reductase and taurine metabolism pathways in SARS-CoV-2 infected hamsters. More interestingly, these sulfur metabolism-related functions were also found to be significantly correlated with *Anaerotignaceae* specific putrescine aminotransferase, which can lead to putrescine degradation and thereby reduced GABA synthesis from microbiome. The significant positive correlations between sulfate reducing bacteria and putrescine aminotransferase may indicate a potential role of these bacteria in the aberrant gut-brain axis and the development of neurological symptoms in COVID-19 patients.

In summary, this study established a reference database as well as a comprehensive workflow based on advanced DIA-MS for hamster metaproteomic study. The application to study the microbiomes in hamsters with SARS-CoV-2 infection demonstrates age- and time-specific alterations of host proteins, microbial taxonomy and functions, as well as their cross-talks. Given the prominent role of the microbiome in diseases and the well-recognized suitability of the hamster model for research on emerging and re-emerging high-consequence infectious diseases, the methodology developed in this study offers a valuable framework for investigating microbiome-related disease or therapeutic mechanisms. Altogether, the hamster microbiome protein databases and the tailored metaproteomic workflow are significant contributions to the disease and drug research with hamsters as animal models, enabling the opportunity to examine the associations of gut microbiota composition and functions with the host responses.

## Methods

### Animal experiment and sample collection

This animal study was conducted in the biosafety level 3 laboratory (BSL3) of the Institut Pasteur de Lille. The protocols were validated by the local committee for the evaluation of the biological risks and complied with current national and institutional regulations and ethical guidelines (Institut Pasteur de Lille/B59-350009). The experimental protocols using animals were approved by the institutional ethical committee “Comité d’Ethique en Experimentation Animale (CEEA) 75, Nord Pas-de-Calais”. The animal study was authorized by the “Education, Research and and Innovation Ministry” under registration number APAFIS#25041-2020040917227851v3. Young (2-month-old, N = 6) and old (22-month-old, N = 6) male Syrian golden hamsters (*Mesocricetus auratus*), equivalent to young adults (∼20 years old) and aged humans (∼80 years old), respectively, were purchased from Janvier Laboratory (Le Genest-Saint-Isle, France). Animals were infected by the intranasal route with 100 µl of DMEM containing 2×10^4^ TCID_50_ (50% of the tissue culture infectious dose) of SARS-CoV-2 (hCoV-19_IPL_France strain)^10, 29^. Feces were collected on day 0, 7, 15, 30 and 45, and stored in -20 °C until processing.

### Protein extraction, trypsin digestion and desalting

#### Stool pre-processing

Enrichment of bacterial cells was performed by differential centrifugation as previously described^48^. Briefly, 0.5 mL glass beads and cold phosphate-buffered saline (PBS) per gram of fecal sample were added to the samples, followed by vortexing and centrifugation at 300 g at 4°C for 5 min to collect the supernatant while the pellets were extracted two more times with cold PBS. The supernatant was further clarified by three centrifugations at 300 g at 4°C for 5 min, then spun down at 14,000 g at 4°C for 20 min to collect the microbial pellet, the pellet was washed twice with cold PBS by resuspending and centrifuging at 14,000 g at 4 °C for 20 min, then frozen until use.

#### Sample lysis

Samples were lysed by resuspending frozen differential centrifuged fecal pellet in 200 µL lysis buffer, consisting of 4% (w/v) SDS, 8M urea in 100mM TEAB. Sample lysates were sonicated using a Bioruptor® Plus sonication device (Diagenode Cat# B01020001) at 50% amplitude, 10 seconds pulse on/off cycle, for 20 min active sonication time at 8°C. The lysate was then centrifuged at 16,000g for 10 min at 8°C to remove any non-lysed debris. Proteins underwent an acetone precipitation by adding 6 volumes of ice-cold acetone, mixing by inversion and incubating at –20°C overnight. Samples were spun down at 16000g at 4°C for 25 min and supernatant was discarded, the remaining pellet was washed twice with 1mL ice-cold acetone. After the last wash and brief air-dry, the protein pellets were stored for further processing as detailed below.

#### In-solution trypsin digestion

The protein pellets were resuspended in 100 µL 0.5% SDS, 8M urea in 100mM TEAB. Protein concentrations were determined using the Pierce BCA Protein Assay Kit (Thermo Fisher Scientific, Cat #23225) following the manufacturer’s protocol using BSA to generate the standard curve. For each sample 100 µg of protein lysate was reduced with 10 mM DTT incubated at 500 rpm at 56°C for 90 min and alkylated with 20 mM IAA for 30 min at room temperature protected from light followed by quenching the alkylation with a further 20 mM DTT. SDS was removed with acetone precipitation and wash as described above. After brief air-dry, the protein pellets were resuspended in 100 µL 0.6M urea in 100mM TEAB buffer and 4 µg trypsin (Promega, Cat#V511B) was added for digestion at 1:25 enzyme:protein at 500 rpm at 37°C overnight. The reaction was stopped by adding 10 µL 10% formic acid to acidify the samples to pH 2-3.

#### Desalting

Desalting was done using C18 columns (Waters Sep-Pak C18 cat # 186002318) in vacuum manifold with all steps performed at 5 psi. The C18 plate was conditioned twice with 200 µl 50% ACN then equilibrated three times with 200 µl 5% ACN / 0.5% TFA. Samples were loaded to the plate then the plate was washed twice 5% ACN / 0.5% TFA and three times with 0.1% FA (v/v). Desalted peptides were then eluted twice with 100 µl 75% (v/v) ACN / 0.1% (v/v) FA. The eluates containing desalted tryptic peptides were then dried on a centrivap (Labconco, Cat#7810010), resuspended in 30 µl 0.1%FA, the concentration determined with Colorimetric Peptide Assay (Pierce, cat # 23275) according to the manufacturer’s directions and samples were diluted to 2 µg/µl with 0.1% FA.

#### Pooling samples and fractionation

Four pooled samples representing four biological categories (Young Pre-infection, Old Pre-infection, Young Post-infection, Old Post-infection) were created by combining samples as follows: Pool 1 - 10 µl each of Young Day 0; Pool 2 -10ul each of Old Day 0; Pool 3 - 2 µl each of Young 7, 15, 30, 45, Pool 4 - 2 µl each of Old Day 7, 15, 30, 45. These pools were fractionated into 8 fractions each using High pH Reversed Phase fractionation kit (Pierce cat # 84868) according to the manufacturer’s directions. Fractions were dried on a centrivap and resuspended in 15 µl 0.1% FA for LC-MSMS analysis.

### LC-MSMS analysis

MS analysis was done using a timsTOF Pro 2 mass spectrometer (Bruker Daltonik, Bremen, Germany) coupled to a nanoElute 2 UPLC system (Bruker Daltonik). The instrument was calibrated prior to analysis with Chip Cube High Mass Reference Standard (Agilent, G1982-85001). A two-column system of HPLC was used consisting of a C8 trap column before separating on a PepSep Twenty-five analytical column (25 cm x 75 µm column packed with 1.9 µm C18 particles) (Bruker Daltonik). Chromatographic separation was achieved at a flow rate of 0.5 µl/min over 48 min in linear steps as follows (solvent A was 0.1% formic acid in water, solvent B was 0.1% formic acid in acetonitrile): initial, 2%B; 40 min, 35%B; 40.5 min, 95%B; 45 min, 95%B; 48 min 95%B. The eluting peptides were analyzed in either DDA-PASEF mode or DIA-PASEF mode in the timsTOFPro 2 mass spectrometer.

#### DDA-PASEF mode

A MS survey scan of 100-1700m/z and ion mobility range of 0.85-1.30Vs/cm2 was performed in the timsTOF MS. During the MS/MS scan, 4 PASEF ramps were run with an intensity threshold of 2500, target intensity of 20000 and a maximum precursor charge of 5. The TIMS analyzer was operated in a 100% duty cycle with equal accumulation and ramp times of 100 ms each and a total cycle tile of 0.53 s. The collision energy was ramped linearly as a function of mobility from 59 eV at LJ 1/K0 = 1.6 Vs/cm2 to 20 eV at 1/K0 = 0.6 Vs/cm2.

#### DIA-PASEF mode

A MS survey scan of 100-1700m/z and ion mobility range of 0.6-1.60Vs/cm2 was performed. The TIMS analyzer was operated in a 100% duty cycle with equal accumulation and ramp times of 100 ms each and a total cycle time estimate 1.8 sec. During DIA-PASEF MS/MS scan, precursors with m/z between 400 and 1200 were defined in 16 scans containing 32 ion mobility steps with an isolation window of 26 Da in each step with 1 Da overlap with neighbouring windows. The collision energy for DIA-PASEF scan was ramped linearly from 59 eV at 1/k0 = 1.3 V·s/cm2 to 20 eV at 1/k0 = 0.85 V·s/cm2.

### Metagenomic Database Generation

This study utilized an in-house shotgun metagenomic sequencing dataset from our companion study with an independent cohort of 3 young (2 months old) and 6 old hamsters (22 months old) for the establishment of a reference protein database for metaproteomics. Details on fecal sample collections, DNA extraction, library generation and sequencing were described in the study by Rodrigues et al.^28^

Low quality sequences were first removed with fastp (version 0.23.4)^49^ with default parameters and host sequences were removed with Kneaddata workflow (https://huttenhower.sph.harvard.edu/kneaddata/) with Bowtie 2^50^ and the hamster genome as reference (GCF_017639785.1_BCM_Maur_2.0). The high quality sequences were then subjected for assembly, gene prediction, taxonomic annotation and quantification, and binning with a previously published SqueezeMeta (v1.6.4) workflow^23^. Briefly, the sequence assembly was performed using Megahit, retaining contigs with >200 bpd. PRINSEQ (0.20.4 lite) was used to quality control for the contigs^51^. tRNA and ribosomal RNA sequences were removed with Aragorn^52^ and Barrnap^53^, respectively, prior to gene prediction with Prodigal^25^. Contig abundances were used for quantitative taxonomic analysis. Taxonomic annotation of contigs was performed by DIAMOND^54^ against the Genbank nr database. Top abundant microbial species were selected based on the calculated taxonomic abundances (MGtax1000 for species with abundance >0.1%, and MGtax10000 for abundance >0.01%). To improve the coverage of the gene catalog database, representative proteome sets were downloaded from UniprotKB Proteomes (45 proteomes for MGtax1000, and 111 proteomes for MGtax10000; Supplementary Table S2) and each was concatenated with gene catalog databases for further evaluations.

To generate a more comprehensive gene catalog database, we downloaded a hamster shotgun metagenomic dataset consisting of 90 metagenomes previously published by Shen et al.^26^ and performed a co-assembly for gene prediction with the same workflow as described above. High quality contigs were further used to construct draft genomes using Metabat2^55^. A total of 926 bins were obtained, and GTDB-Tk (v2.4.0)^56^ was used for taxonomic annotation. Among these, 706 bins were annotated as bacteria, with 251 classified at the species level and 423 at the genus level. Whole genomes for the 251 species were downloaded from GenBank and combined with the assembled bins. The 706 annotated genomes, along with the 251 downloaded genomes, were then digested in silico using Rapid Peptides Generator (RPG) v2.0.5 with trypsin^57^. After filtering out peptides with lengths outside the range of 7-30 amino acids, a total of 82,350,794 peptides were retained for database construction for MetaX analysis. The workflow was illustrated in Supplementary Figure 15.

### Reduced protein database generation with DDA dataset

To generate a reduced protein database, DDA-PASEF data of pooled fractions were searched against the MGDB-V2 database with MSFragger (version 4.0)^58^ implemented in FragPipe (v21.1). The default workflow was used with the following changes: a database split factor of 20, no mass calibration and parameter optimization for MSFragger search, and spectral library generation with ciRT for RT calibration. An FDR threshold of 1% was used for both peptide and protein identifications. The sequences of all identified proteins from the MSfragger search, including indistinguishable proteins, were extracted from original MGDB-V2 using an in-house Perl script to generate a reduced FASTA database.

### DIA-NN search for identification and quantitation of DIA-MS dataset

DIA-PASEF data were processed with DIA-NN (v1.9.1)^59^ in library-free mode using the reduced FASTA database generated through MSFragger searches as described above. To identify and quantify host proteins we appended the Golden hamster proteome (*Mesocricetus auratus* with 20389 entries; downloaded from UniProtKB on July 24th, 2024) to the reduced database. Default settings were used for the DIA-NN search with a precursor and protein level FDR threshold of 1%. MaxLFQ intensities of the identified proteins or peptides were used for all the downstream data analysis.

### Downstream taxonomic and functional analysis

Taxonomic annotation and analysis were performed using MetaLab (v2.3.2)^60^ with the peptide data matrix (report.pr_matrix.tsv) as input and UniPept for taxonomic assignment^61^. A minimum of 3 distinctive peptides was required for confident identification of taxa. The taxon intensity was calculated with sum intensity of all distinctive peptides assigned to it and then normalized at each taxonomic rank level to obtain relative abundances for further statistical analysis.

For microbiome functional analysis, all host proteins were first excluded from the quantitative data matrix (report.pg_matrix.tsv) and the remaining microbiome protein abundances were then normalized for further analysis. Functional annotation of microbial proteins was performed using GhostKOALA with default parameters^32^. The KEGG GENES database file was set as “genus_prokaryotes + family_eukaryotes + viruses”. The normalized intensities of microbial proteins annotated with the same KO were summed up to obtain KO abundance for further statistical analysis.

### Statistical data analysis and visualization

In this study, we kept proteins or microbial functions with valid non-zero values in at least 70% samples for statistical analysis, including principal component analysis (PCA) and partial least-squares-discriminant analysis (PLS-DA). The intensities were log2-transformed and the missing values were imputed with k-nearest neighbor (KNN) algorithm when needed. PCA was performed in R with the princomp function and visualized with ggplot2. PLS-DA was performed in MetaboAnalyst 5.0^62^ to calculate the variable importance in projection (VIP) of each protein. VIP is a normalized value, and a VIP > 1 indicates a significant contribution to the PLS-DA model. Hierarchical clustering and heatmap of identified VIP proteins or functions were performed using the R package ComplexHeatmap^63^.

Statistical analysis of taxonomic composition was performed using MaAsLin2 (Galaxy Version 1.8.0)^37^. Young-Day7 and Old-Day7 groups were set as fixed effects and the others were set as reference group. The abundance data were log transformed prior to analysis using the LM (linear model) method. Statistical P values were corrected with Benjamini-Hochberg method and a corrected p-value <0.05 was deemed as significant in this study.

Diagrams in Figures 1 and 2 were created in BioRender (Zhang, X. (2023) BioRender.com/f82w697). Co-occurrence analysis was performed using iMetaShiny^64^ (https://shiny.imetalab.ca/) and visualized using Cytoscape (v3.10.2)^65^. The sankey plot was generated using SankeyMATIC (https://sankeymatic.com/).

## Data availability

All MS proteomics data that support the findings of this study have been deposited to the ProteomeXchange Consortium (http://www.proteomexchange.org). Raw metagenomic sequencing data was submitted to NCBI sequence read archive (BioProject: PRJNA1121382). The metagenomics derived hamster gut microbial gene catalog databases along with all FASTA files of binned genomes, annotations, and the MetaX database, have been uploaded to Zenodo (https://zenodo.org/records/13909289).

## Acknowledgements

This work was supported by funding from the Government of Canada through Health Canada. Q.W. is supported by a stipend from the TECHNOMISE NSERC-CREATE program. We gratefully acknowledge Drs. Simon Sauvé, Michael Rosu-Myles, Michael Johnston and Julie Shay from Health Canada for their critical comments and edits to this manuscript.

## Notes

### Competing Interest Statement

The authors have declared no competing interest.

