## Supplemental materials for "Metaproteomics reveals age-specific alterations of gut microbiome in hamsters with SARS-CoV-2 infection"

Marybeth Creskey, *et al*.

**This supplementary information includes:**

- Supplementary Table S1-9
- Supplementary Figure 1-15

**Supplementary Tables:**

**Supplementary Table S1**. Top abundant hamster fecal microbial taxa identified with in-house metagenomics sequencing data. (.xlsx)

**Supplementary Table S2**. Top abundant species and representative UniprotKB proteomes for generating MGtax databases. (.xlsx)

**Supplementary Table S3**. VIP1 host proteins differentiating Young-Day7, Old-Day7 and others. (.xlsx)

**Supplementary Table S4**. VIP1 host proteins differentiating recovery phase hamsters with baseline. (.xlsx)

**Supplementary Table S5**. VIP1 microbial KOs differentiating Young-Day7, Old-Day7 and others. (.xlsx)

**Supplementary Table S6**. VIP1 microbial KOs differentiating recovery phase hamsters with baseline. (.xlsx)

**Supplementary Table S7**. Identified taxa and corresponding distinctive peptide numbers with metaproteomics. (.xlsx)

**Supplementary Table S8**. Significantly changed taxa in Young-Day7 and Old-Day7 groups identified by MaAsLin2. (.xlsx)

**Supplementary Table S9**. Significantly changed taxa in recovery phase groups identified by MaAsLin2. (.xlsx)

**Supplementary Figures:**

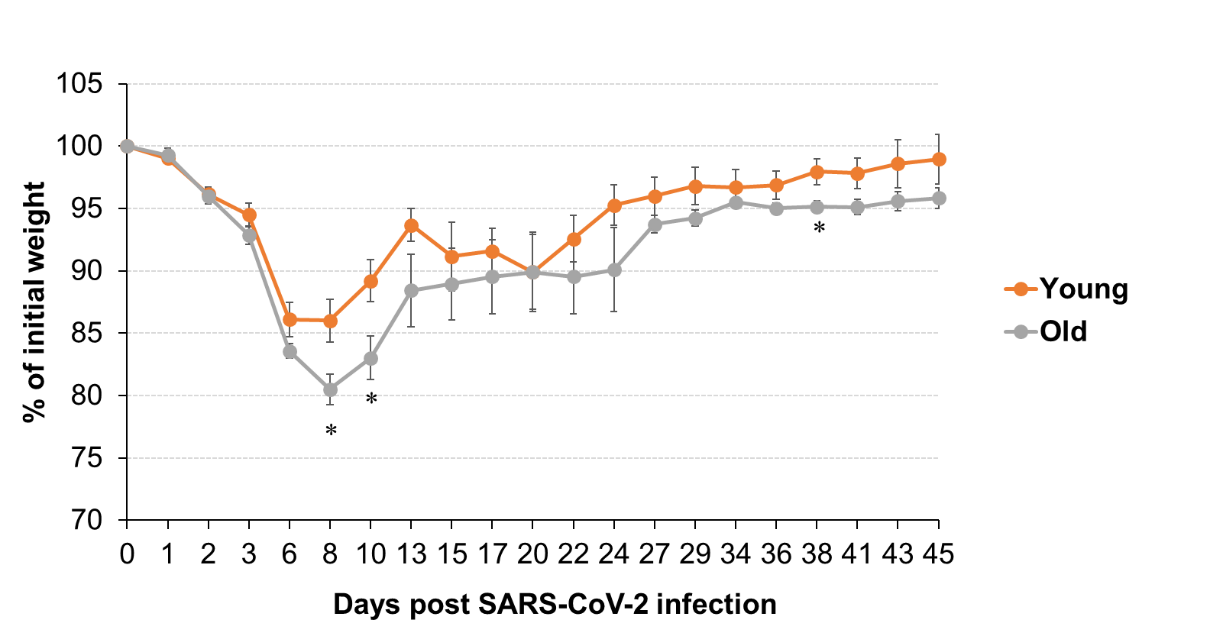

**Supplementary Figure 1**. Percentage of body weight changes of young and old hamsters with SARS-CoV-2 infections. Statistical significance test was performed between young and old groups at each time point with t-test; * p < 0.05.

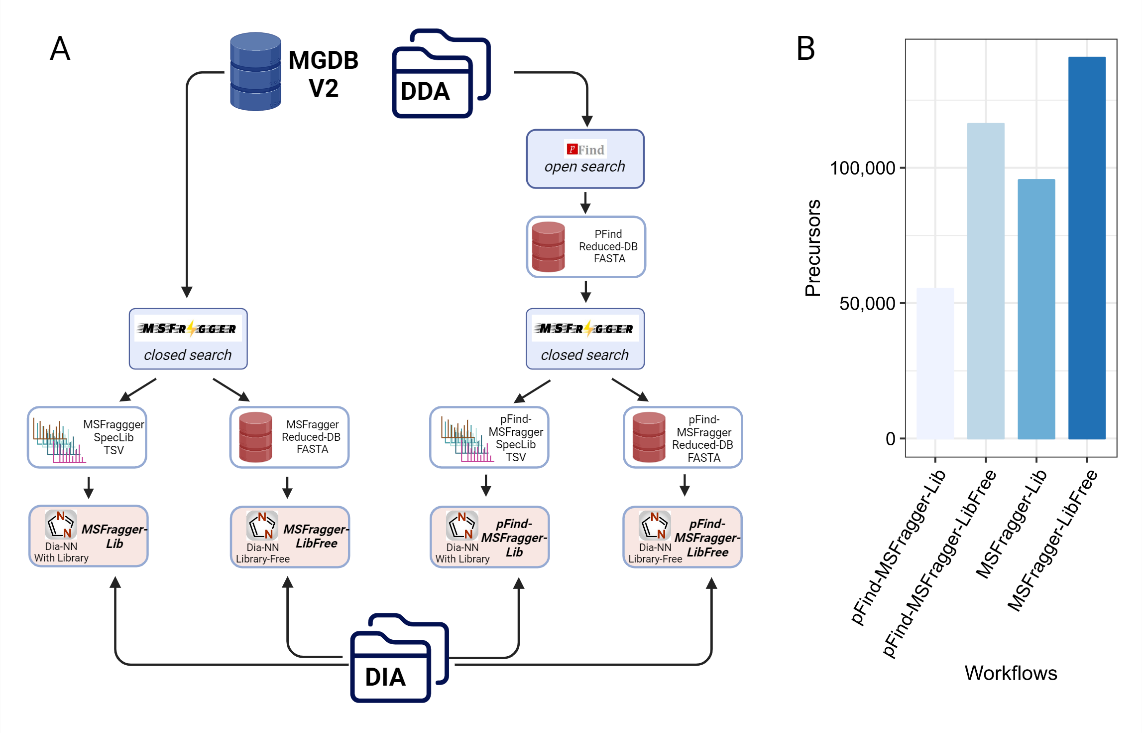

**Supplementary Figure 2**. Evaluation of different bioinformatic workflows to generate databases for DIA-MS data analysis. Numbers of total precursors identified with each workflow are shown.

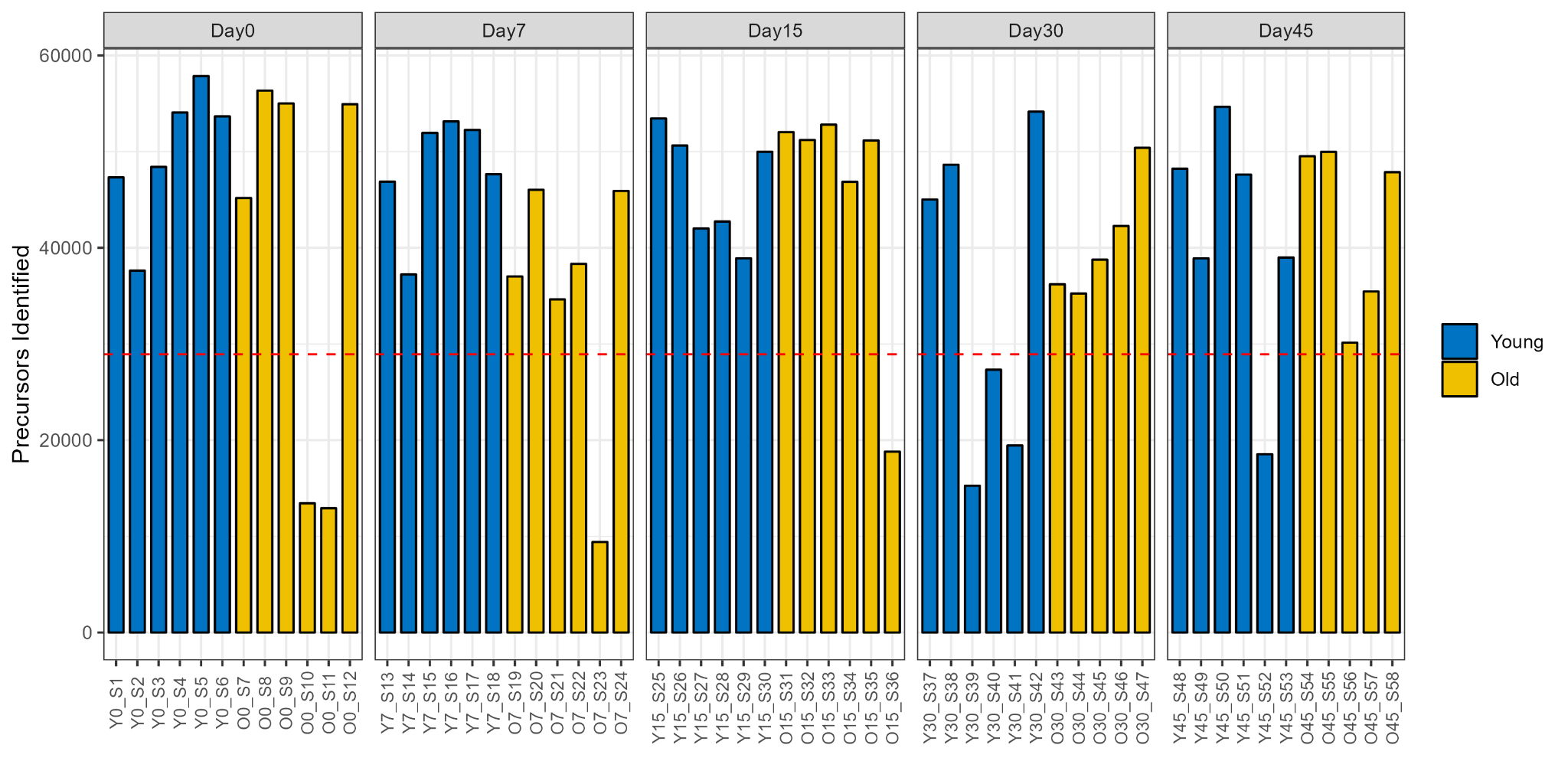

**Supplementary Figure 3**. Precursor identification of each sample with DIA-NN. Red line in the plot indicates the half of the maximum number of identification. The 8 samples with precursor identification less than this number were removed for further statistical analysis.

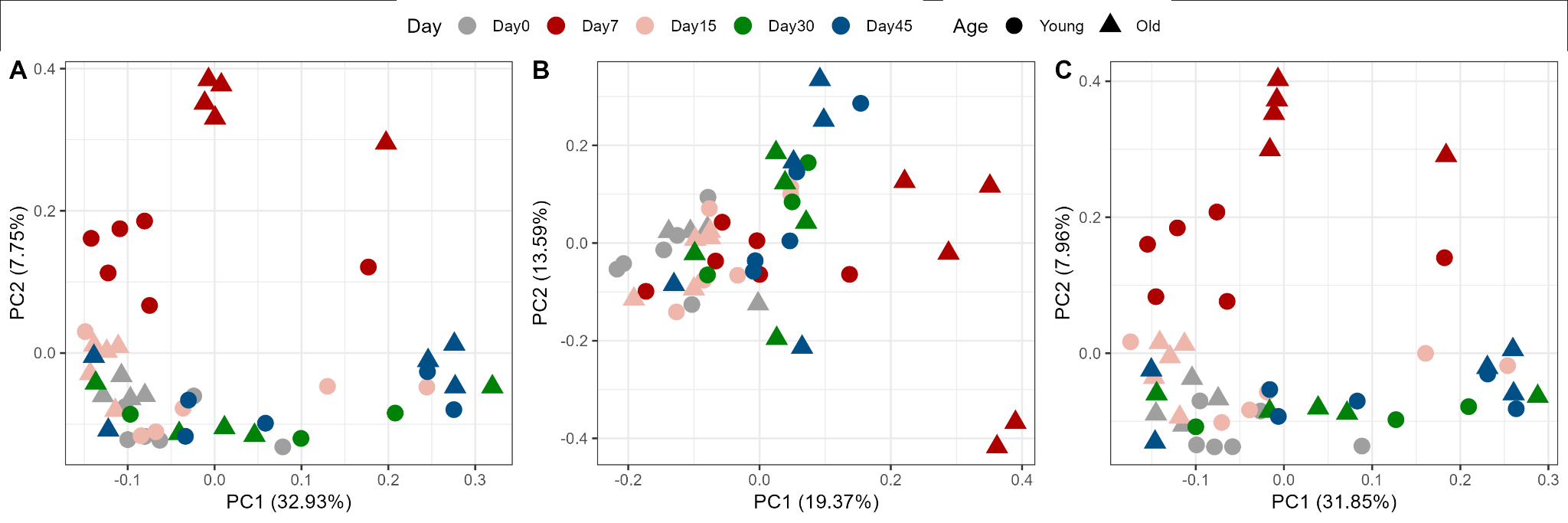

**Supplementary Figure 4**. PCA score plots using quantified microbiome or host proteins (without facet, related to Figure 2). PCA was performed using the normalized and log2-transformed protein intensities for all (A), host proteins only (B) or microbiome proteins only (C). Only proteins that were quantified in >70% of the samples were used for analysis. PCA score plots were generated with R ggplot2.

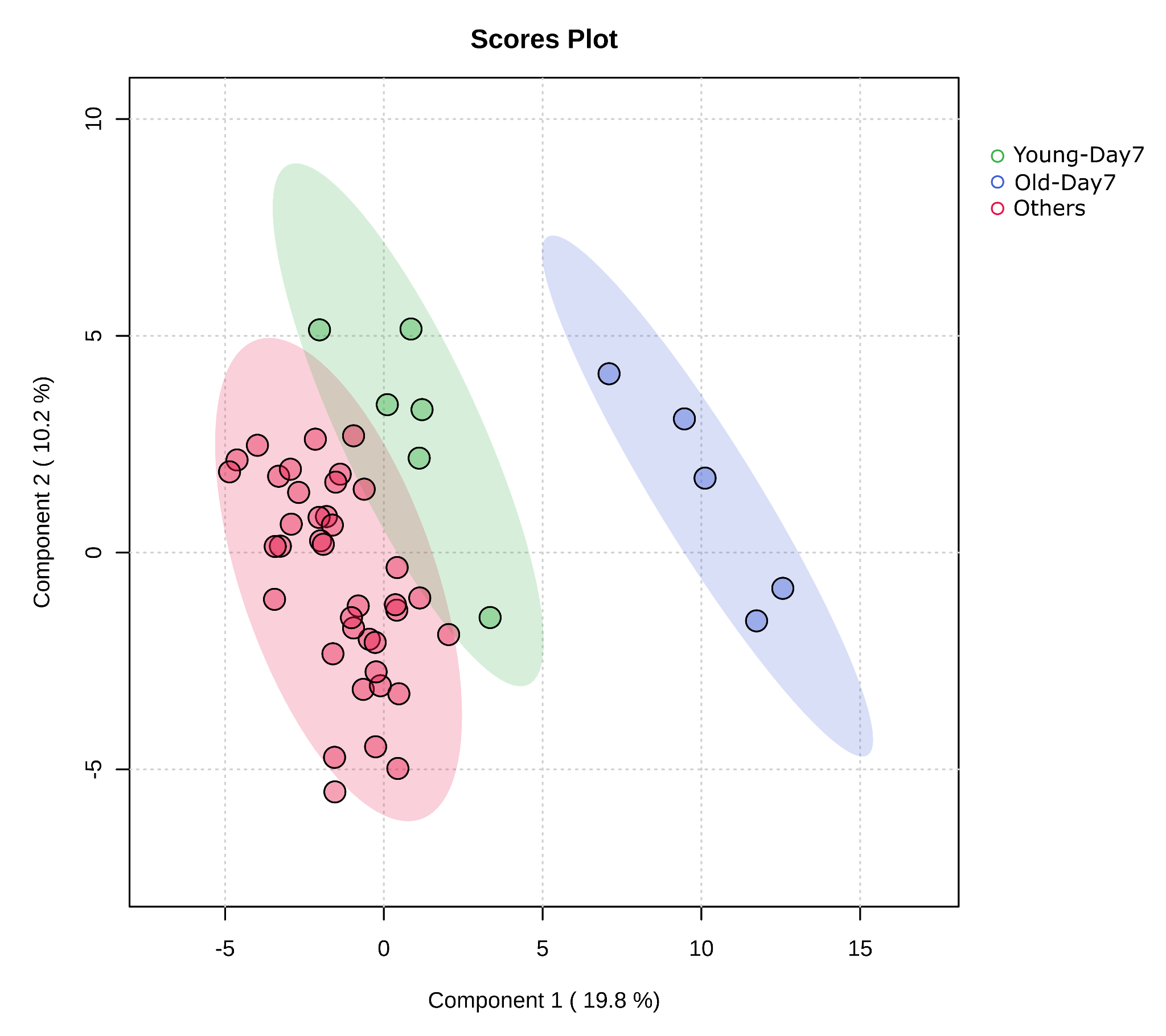

**Supplementary Figure 5**. PLS-DA score plot of quantified host proteins. The normalized abundance data was log2-transformed for PLS-DA analysis. Only proteins that were quantified in >70% of the samples were used for analysis and missing values were imputed with KNN algorithm. PLS-DA was performed using MetaboAnalyst with samples grouped into three clusters: Young-Day7, Old-Day7, and the others.

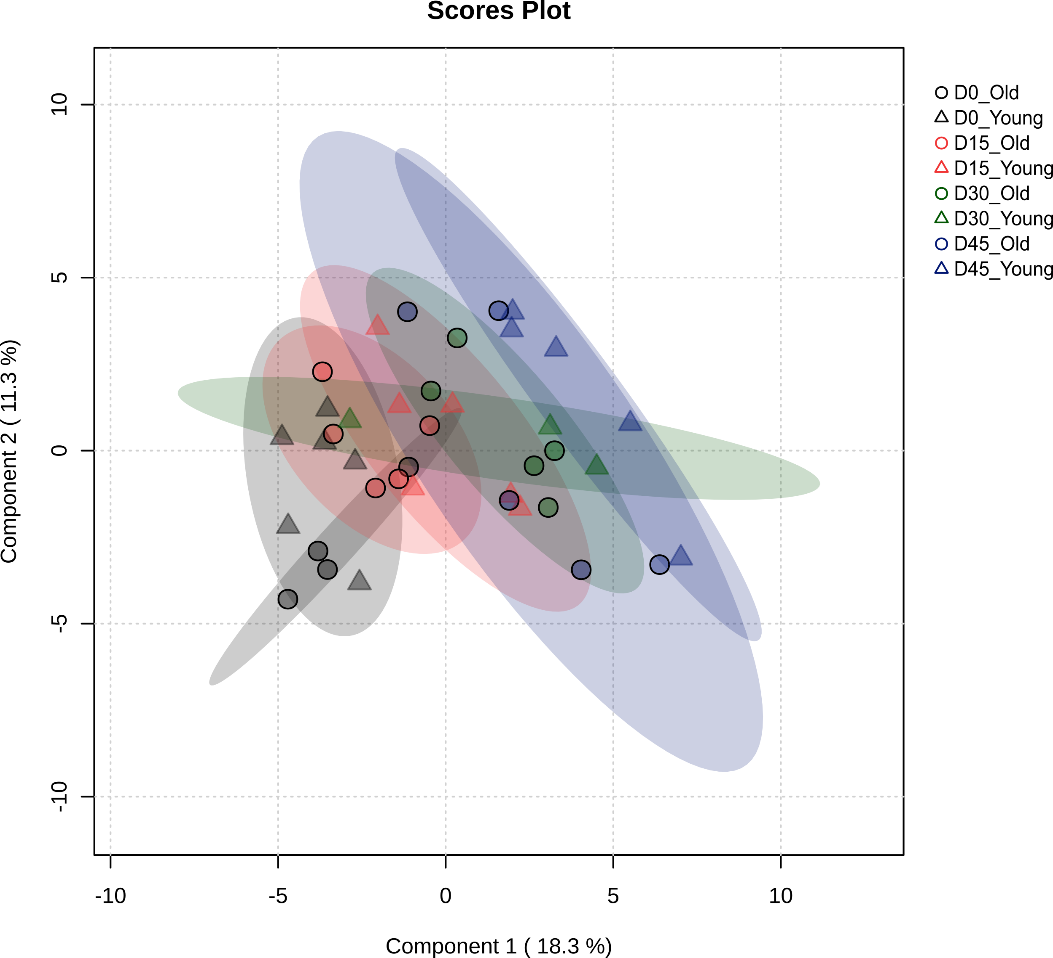

**Supplementary Figure 6.** PLS-DA score plot of quantified host proteins in samples excluding Day 7. The normalized abundance data was log2-transformed for PLS-DA analysis. Only proteins that were quantified in >70% of the samples were used for analysis and missing values were imputed with KNN algorithm. PLS-DA was performed using MetaboAnalyst.

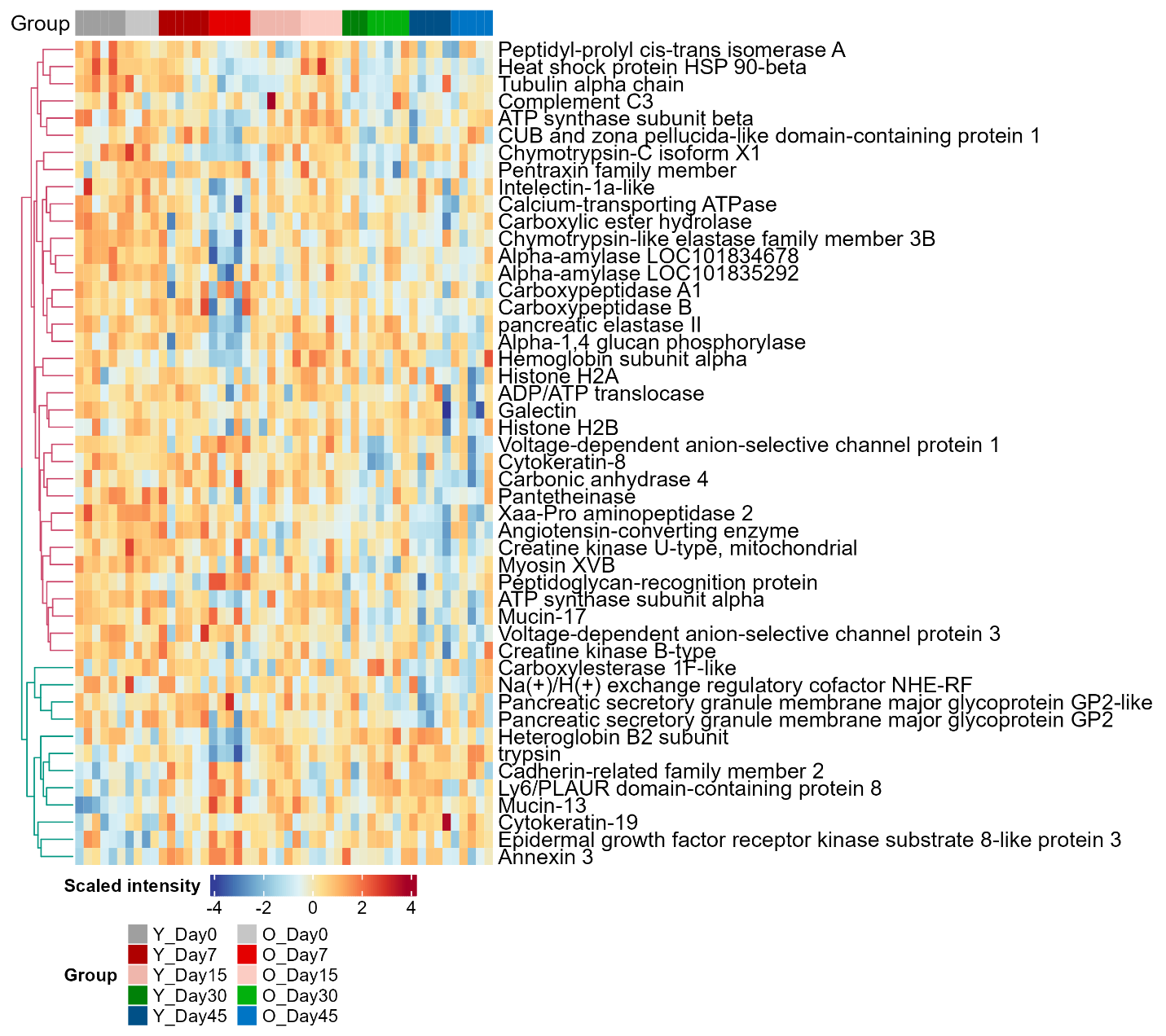

**Supplementary Figure 7.** Heatmap of differentially abundant host proteins in recovery phase. Protein intensities were log2 transformed, scaled and are displayed as colours ranging from blue(low) to red(high) as shown in the key. Heatmap and clustering for rows are performed using the R ComplexHeatmap package. Day 7 samples were excluded for clustering for rows but shown for visualization.

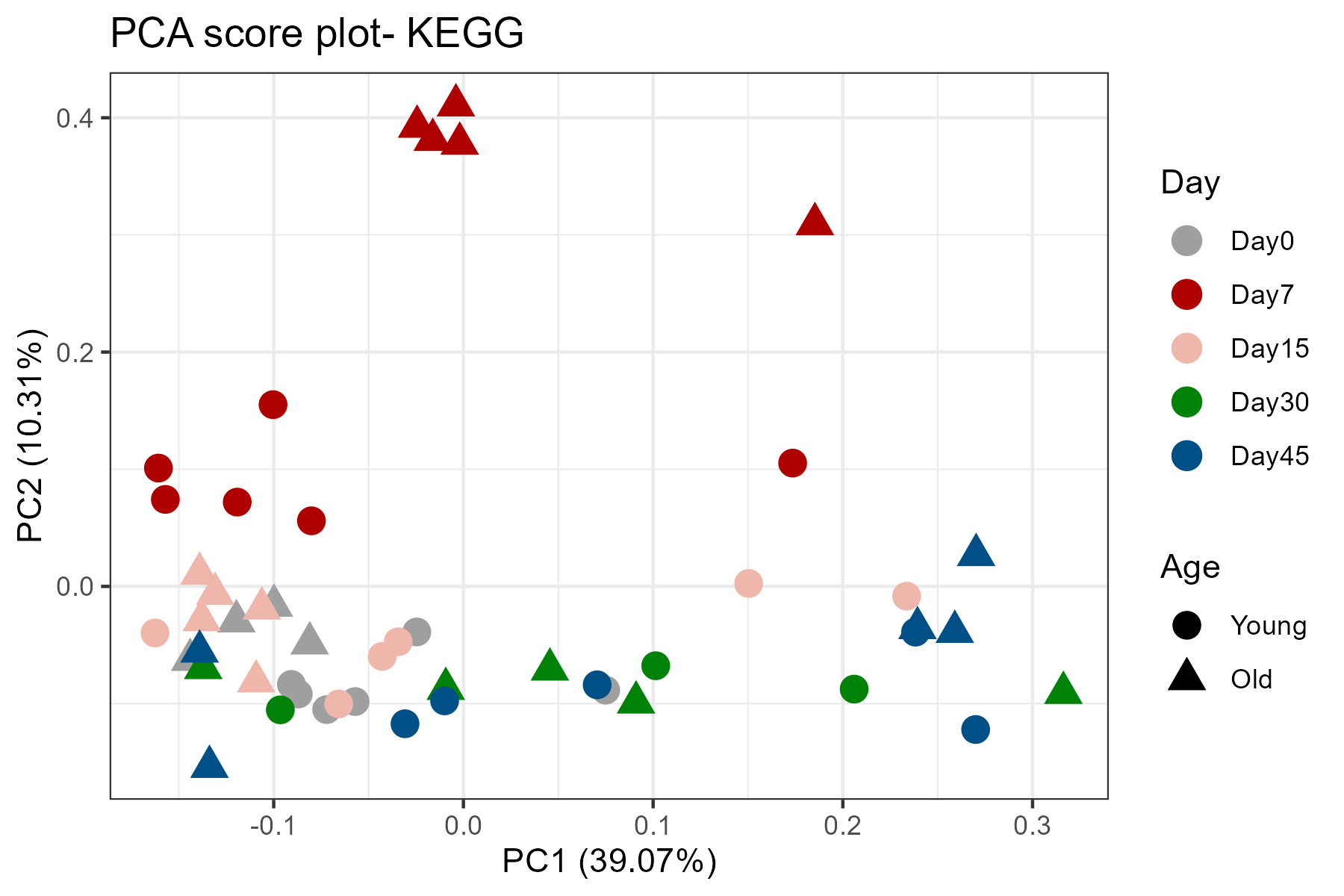

**Supplementary Figure 8**. PCA score plots of quantified KEGG orthologues of microbiome. PCA was performed using the normalized and log2-transformed KO intensities. Only KOs that were quantified in >70% of the samples were used for analysis and missing values were imputed with KNN algorithm. PCA score plots were generated with R ggplot2.

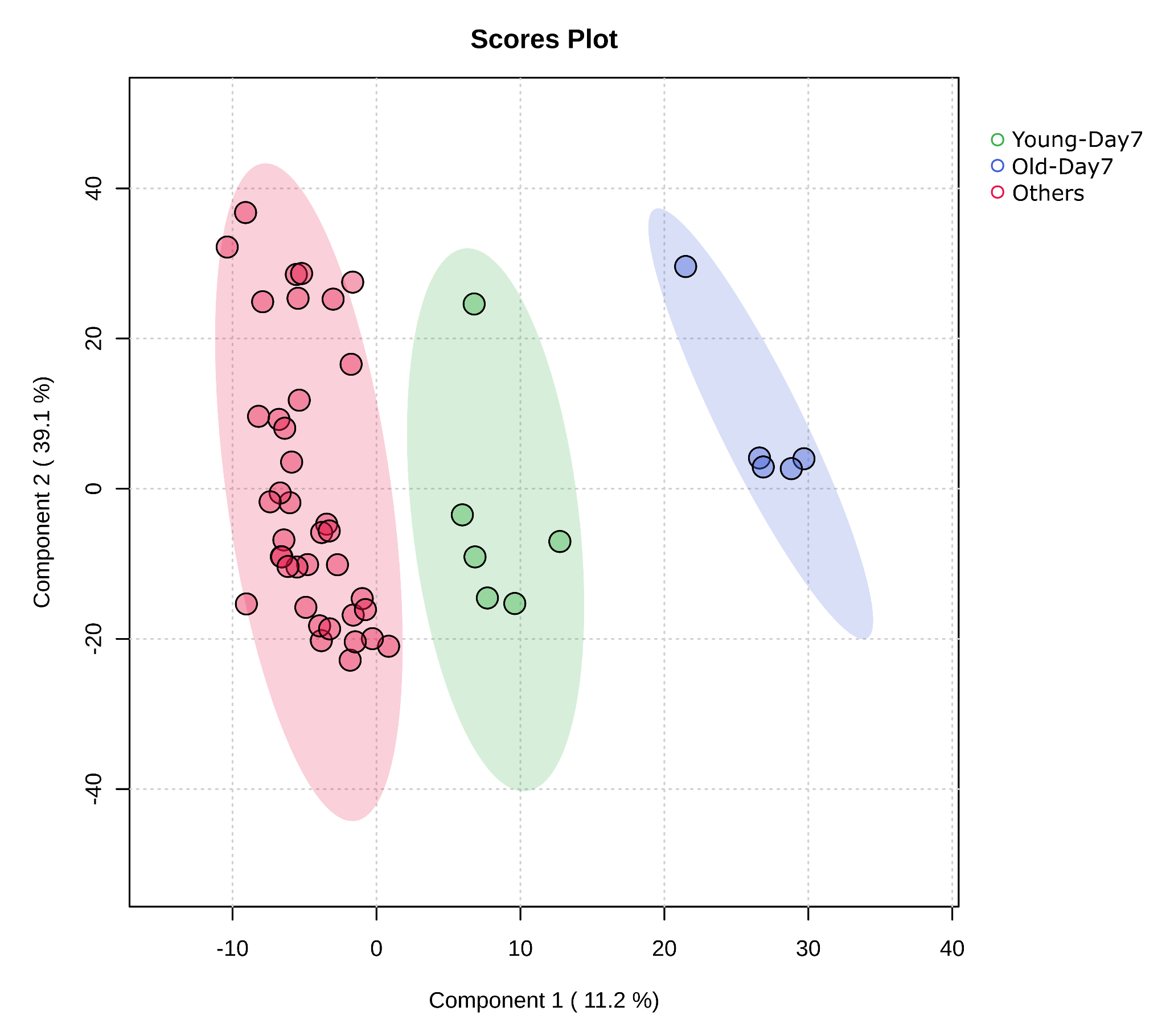

**Supplementary Figure 9**. PLS-DA score plot of quantified KEGG orthologues of microbiome. The normalized abundance data was log2-transformed for PLS-DA analysis. Only KOs that were quantified in >70% of the samples were used for analysis and missing values were imputed with KNN algorithm. PLS-DA was performed using MetaboAnalyst with samples grouped into three clusters: Young-Day7, Old-Day7, and the others.

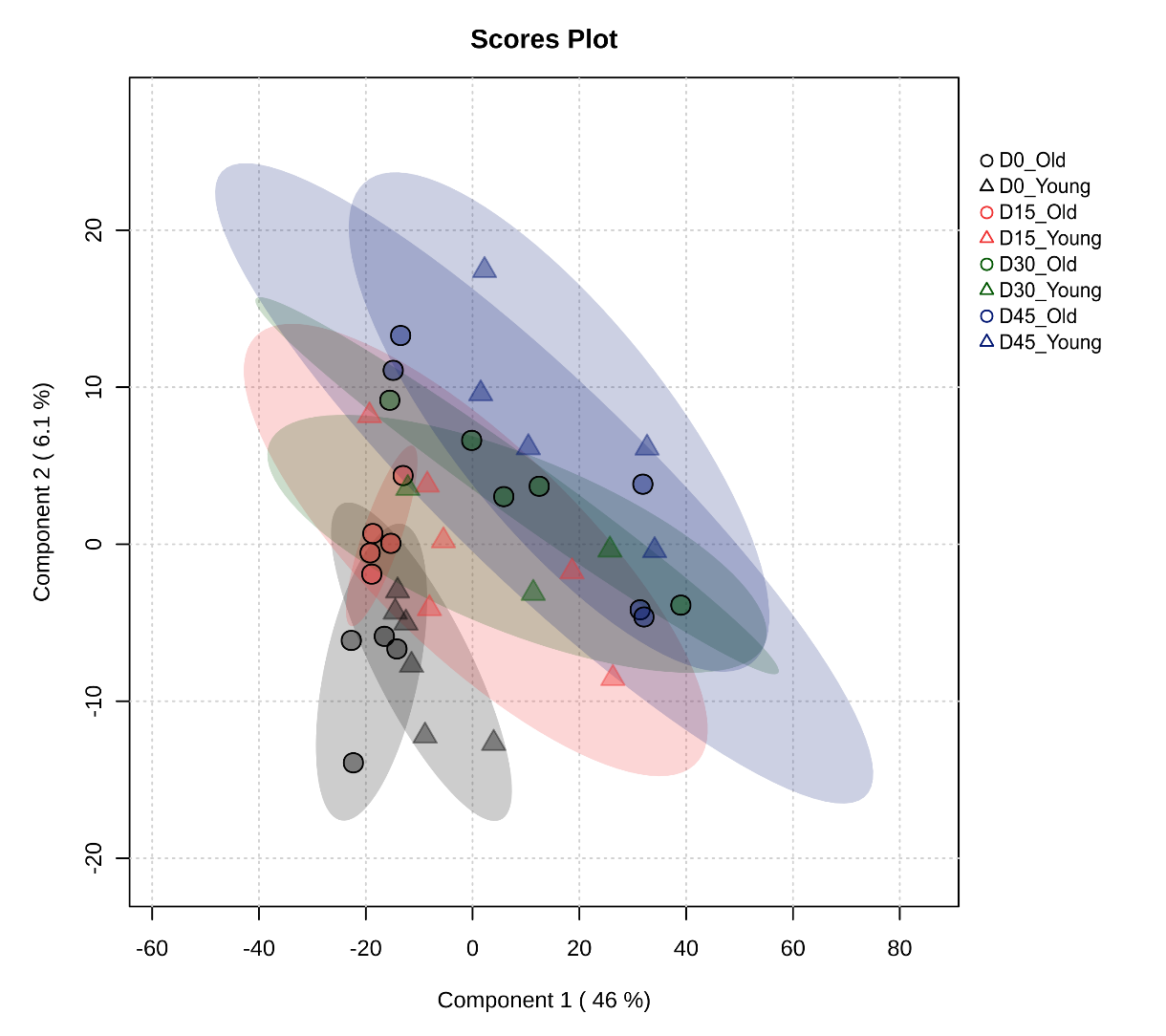

**Supplementary Figure 10.** PLS-DA score plot of quantified KEGG orthologues in samples excluding Day 7. The normalized abundance data was log2-transformed for PLS-DA analysis. Only KOs that were quantified in >70% of the samples were used for analysis and missing values were imputed with KNN algorithm. PLS-DA was performed using MetaboAnalyst.

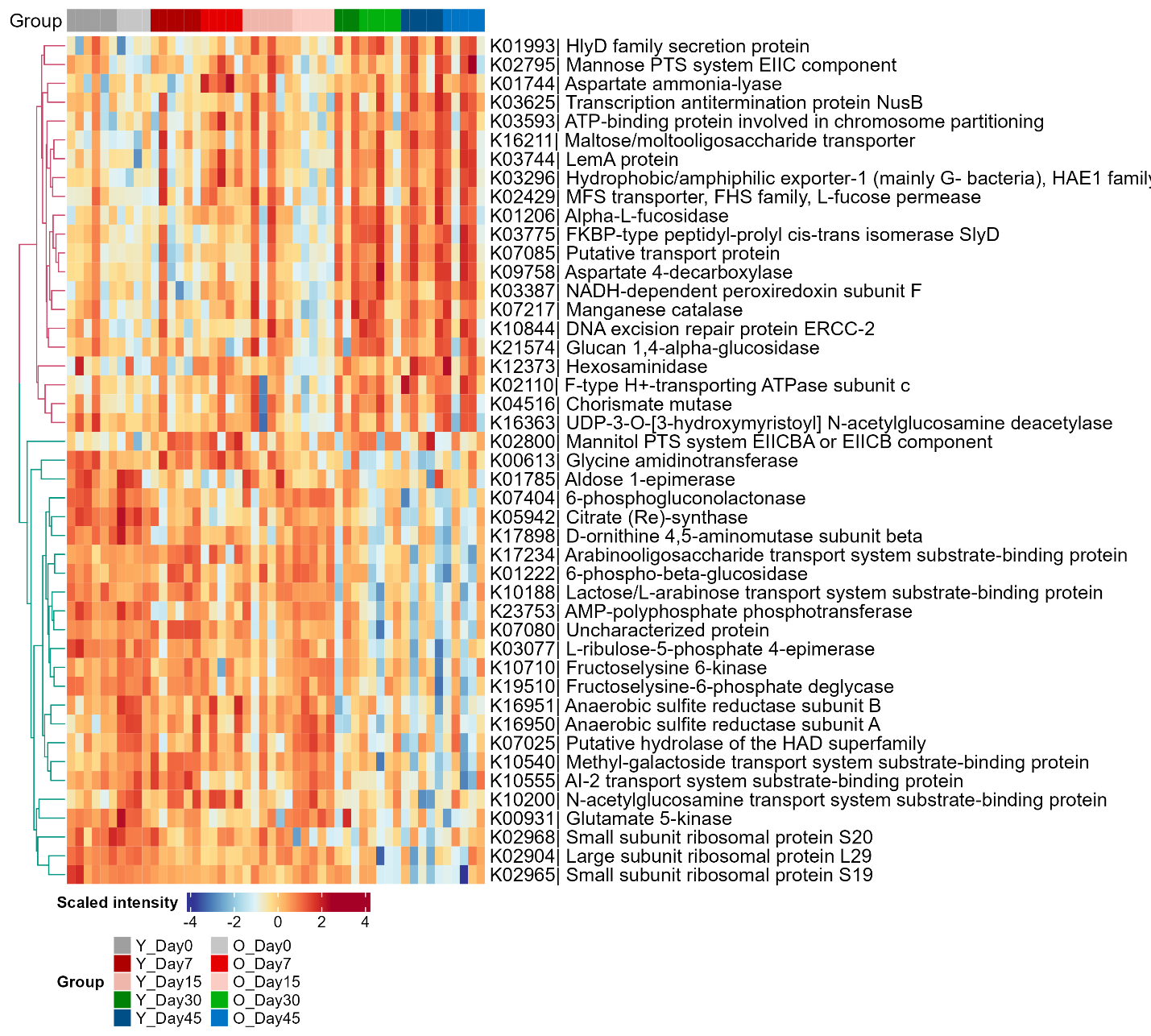

**Supplementary Figure 11.** Heatmap of differentially abundant microbial KOs in recovery phase. KO intensities were log2 transformed, scaled and are displayed as colours ranging from blue(low) to red(high) as shown in the key. Heatmap and clustering for rows are performed using the R ComplexHeatmap package. Day 7 samples were excluded for clustering for rows but shown for visualization.

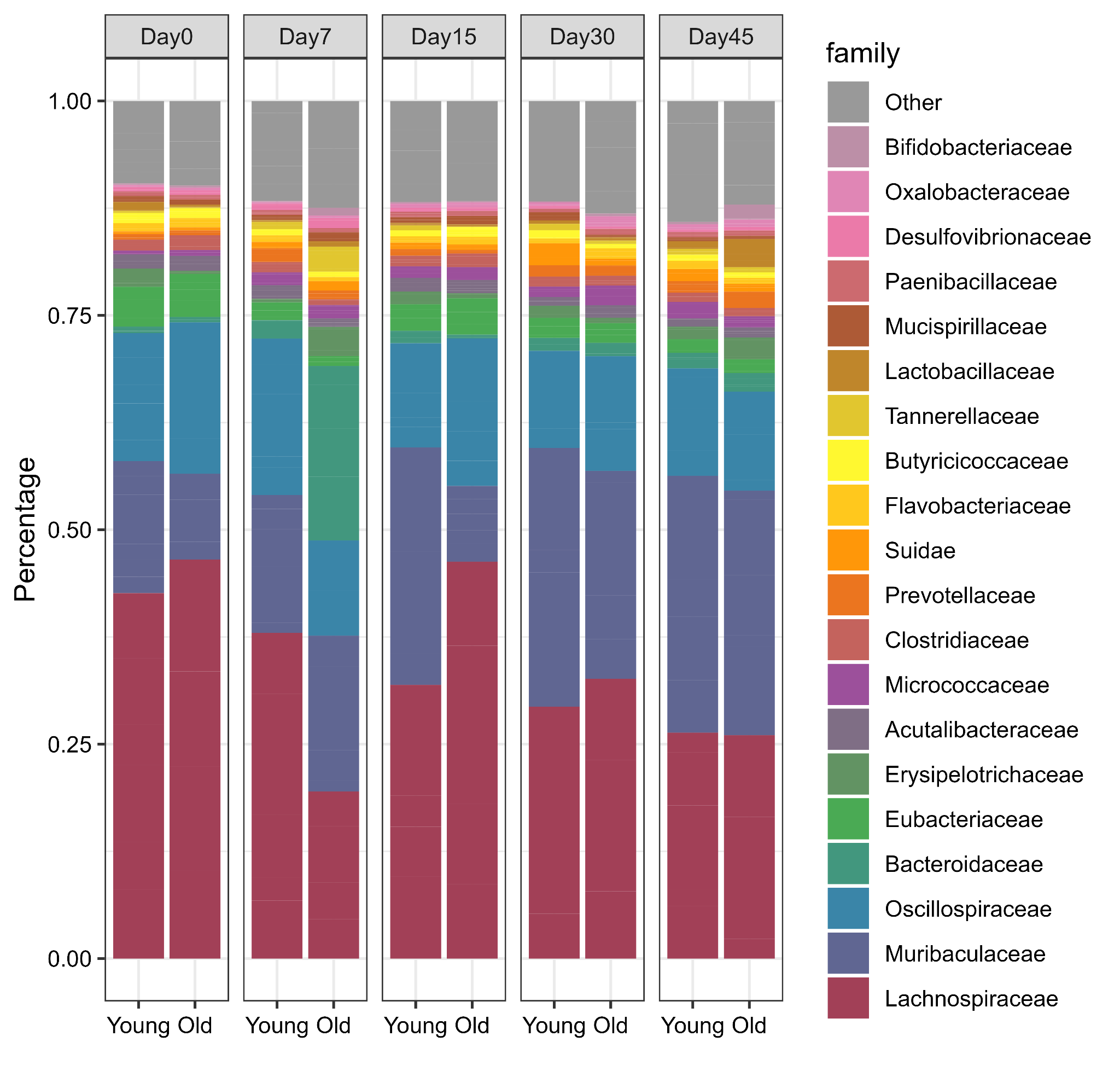

**Supplementary Figure 12**. Family level microbial compositions in hamsters with SARS-CoV2 infection. The most abundant families were shown with remaining families grouped as ‘Other’ (grey) in the plot. Stacked barplot was generated using ggpubr in R.

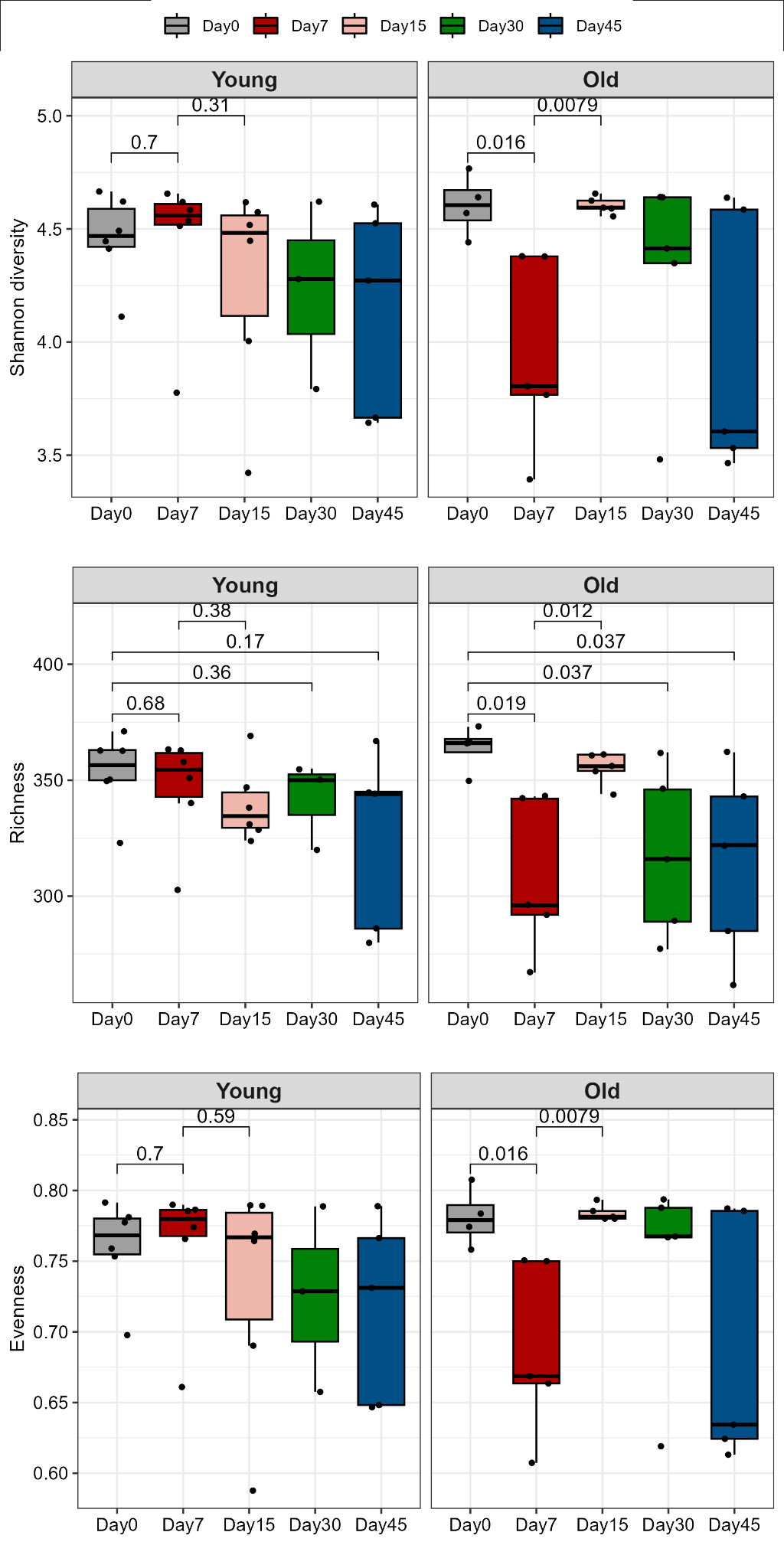

**Supplementary Figure 13**. Species diversity of hamster microbiomes with SARS-CoV2 infection. Diversity, richness and evenness were shown. Pairwise statistical comparisons were performed using the Wilcoxon test for Young and Old groups separately. P values were indicated for comparisons that are significant for either Young or Old groups.

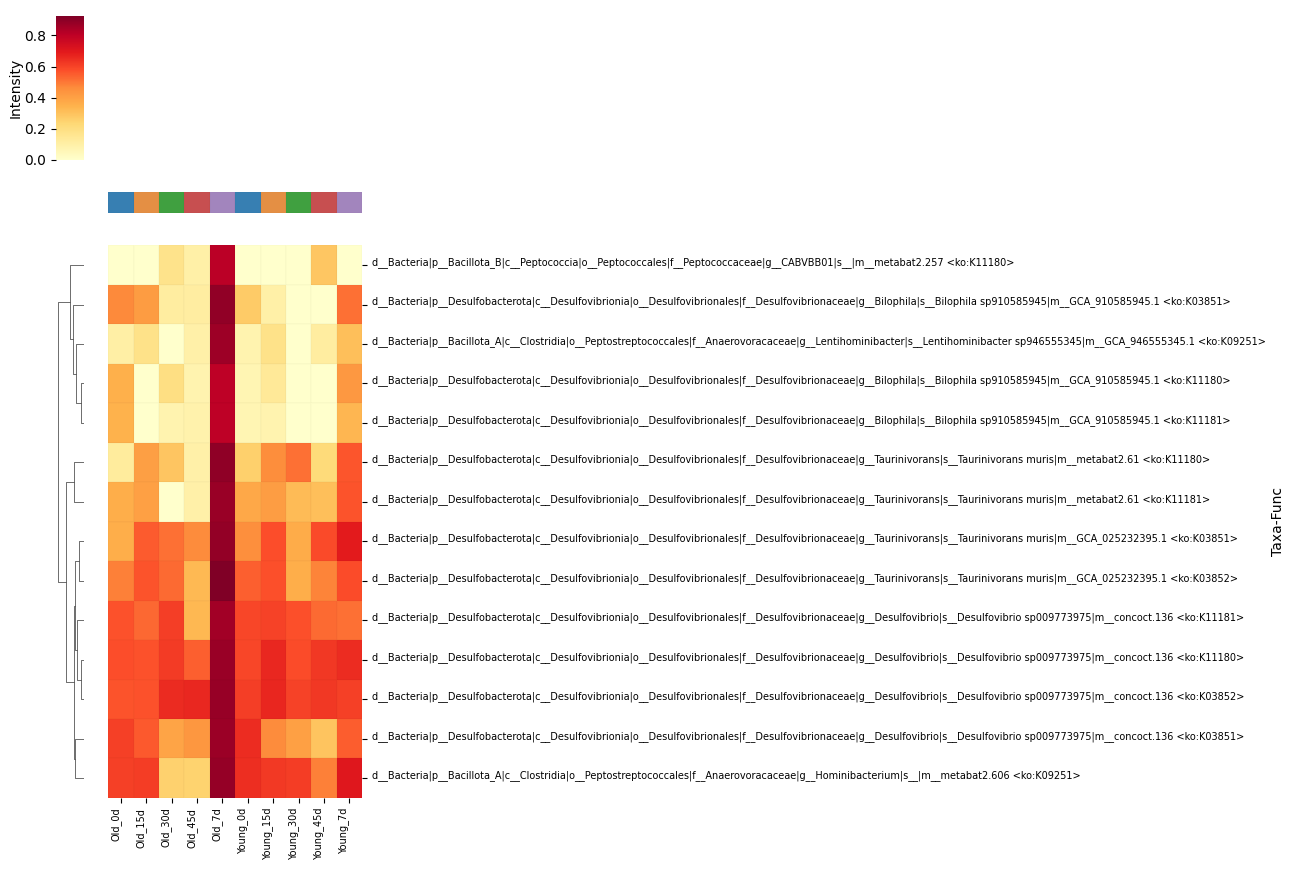

**Supplementary Figure 14**. Heatmap of Operational taxon-function units (OTFs) with the functions implicated in sulfur metabolism in Network B (Figure 8). OTF analysis and heatmap were performed using MetaX (https://github.com/byemaxx/MetaX).

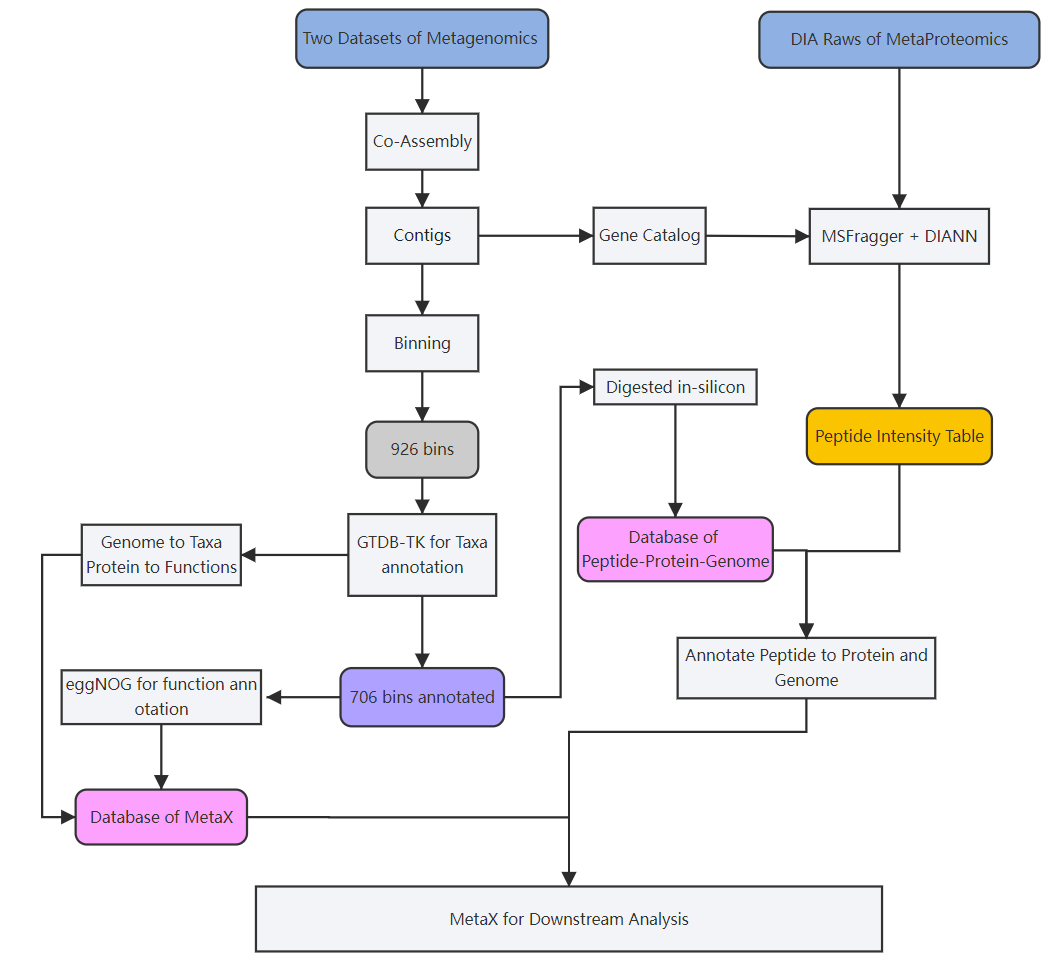

**Supplementary Figure 15**. Bioinformatic workflow to establish database for MetaX analysis.
